# A multi-tiered mechanical mechanism shapes the early neural plate

**DOI:** 10.1101/2023.06.21.545965

**Authors:** Angus Inman, Judith E. Lutton, Elisabeth Spiritosanto, Masazumi Tada, Till Bretschneider, Pierre A. Haas, Michael Smutny

**Affiliations:** Centre for Mechanochemical Cell Biology and Division of Biomedical Sciences, Warwick Medical School, University of Warwick, Coventry CV4 7AL, UK; Department of Computer Science, University of Warwick, Coventry, CV4 7AL, UK; Department of Cell and Developmental Biology, University College London, Gower Street, London, WC1E 6BT, UK; Max Planck Institute for the Physics of Complex Systems, Nöthnitzer Str. 38, 01187 Dresden, Germany; Max Planck Institute of Molecular Cell Biology and Genetics, Pfotenhauerstr. 108, 01307 Dresden, Germany; Center for Systems Biology Dresden, Pfotenhauerstr. 108, 01307 Dresden, Germany

## Abstract

The formation of complex tissues during embryonic development requires an intricate spatiotemporal coordination of local mechanical processes regulating global tissue morphogenesis. Here, we uncover a novel mechanism that mechanically regulates the shape of the anterior neural plate (ANP), a vital forebrain precursor, during zebrafish gastrulation. Combining *in vivo* and *in silico* approaches we reveal that the ANP is shaped by global tissue flows regulated by distinct force generating processes. We show that mesendoderm migration and E-cadherin-dependent differential tissue interactions control distinct flow regimes in the neuroectoderm. Initial opposing flows lead to progressive tissue folding and neuroectoderm internalisation which in turn provide forces driving ANP tissue reshaping. We find that convergent extension is dispensable for internalisation but required for ANP tissue extension. Our results highlight how spatiotemporal regulation and coupling of different mechanical processes between tissues in the embryo controls the first folding event in the developing brain.

## Introduction

Correct shaping and patterning of tissues and organs is a fundamental process during embryonic development which is driven by coordinated cell and tissue level events such as cellular rearrangements and acquisition of specialised cell fates. While biochemical signalling and gene regulation control cell fate specification and tissue patterning, physical forces also play a key role as direct drivers of tissue shape ^1–3^. Notably, mechanical forces regulate local and global morphogenetic events during embryonic development, including tissue movement, shape changes, growth and folding ^4–6^. Understanding the formation of dynamic three-dimensional (3D) tissues remains especially challenging due to the coordination of sequential cellular rearrangements along multiple body axes within the tissue and interactions with cells from neighbouring tissues ^7–10^. Robust tissue morphogenesis requires orchestrated spatiotemporal control of both intrinsic stresses, such as generated by the cellular actomyosin network and extrinsic stresses, including those produced by neighbouring cells or fluid forces ^6, 11^. In particular, coupling of forces between cells within a tissue or between adjacent tissues is crucial to drive large-scale tissue remodelling events, including cell intercalations during convergent extension, tissue-wide cellular flows and tissue folding events ^12–18^. Notably, these large-scale rearrangements do not function in isolation but can generate forces that feedback into tissue morphogenesis and patterning. For example, a force gradient established in response to FGF8 morphogen signalling in the hindgut-forming endoderm has been shown to drive collective movements of non-contractile cells in the chick embryo^19^.

The brain is one of the most complex tissues and undergoes continuous shape changes from the earliest stages of development. In particular, the vertebrate neural plate, a vital precursor of the central nervous system, undergoes extensive rearrangements during neurulation to form the neural tube and nerve cord ^20–22^. Despite extensive research into the mechanisms underlying neural tube formation during neurulation, little is currently known about neural plate morphogenesis during gastrulation ^20, 23, 24^. Failures of functional neural plate formation in the gastrula result in aberrant tissue shape and position and are associated with severe defects of the brain and nervous system in later stages, as evidenced by many mutants identified in *Danio rerio* (zebrafish) ^25–27^. Major efforts have been directed at constructing neural plate fate maps ^28, 29^ and to dissect the various morphogen signalling pathways and downstream transcription factors regulating neural plate regionalisation ^30^. Yet, our understanding about the underlying mechanisms driving cell and tissue morphogenesis during early neural plate formation in the gastrula remains very limited.

Here, we address this question by investigating how the anterior neural plate (ANP), a precursor of the forebrain, is shaped during zebrafish gastrulation. Using a combination of experiments and simulations, we show that ANP cell and tissue morphogenesis is regulated by a spatiotemporally controlled sequence of extrinsic forces and mechanical tissue coupling, and reveal the underlying mechanisms controlling the earliest event of tissue folding in the future forebrain. Our findings highlight the importance of mechanical coordination of distinct morphogenetic events beyond tissue boundaries as a major regulator of shaping complex multi-layered tissues in the developing embryo.

## Results

### The early ANP is dramatically reshaped during late gastrulation

Previous studies suggest that neuroectoderm progenitors show limited cellular rearrangements after regional specification during the early formation of the anterior neural plate (ANP) ^23, 28^. However, ANP tissue shape is not preserved during gastrulation ^31^, which raises the intriguing question of how cell and tissue dynamics are coupled to shape the ANP. To address this problem, we studied ANP tissue dynamics throughout gastrulation by using *Tg(otx2:Venus)* transgenic zebrafish embryos ^32^ that allow visualisation of tissue-specific neuroectoderm progenitor cell fate specification in the prospective ANP (Fig. 1a,b). We observed otx2:Venus expression from ~7.5 hours post fertilisation (hpf) onward, with neuroectoderm progenitor cells initially displaying a scattered distribution and irregular tissue outlines (Supplementary Video 1, Fig. 1b). Analysis of the overall tissue dynamics revealed that little tissue shape changes occurred prior to 8.5 hpf (Fig. 1c–e). However, during later gastrulation stages (8.5–10 hpf) tissue surface area and mediolateral (ML) width decreased rapidly (Fig. 1c,d) while tissue volume was largely preserved (Supplementary Fig. 1a). Notably, ML tissue width of the tissue was reduced sequentially from anterior to posterior (Fig. 1b; Supplementary Fig. 1b) reminiscent of rotational movements leading to bending of the anterior-lateral tissue edges towards the dorsal midline. This tissue rearrangement culminated in an arrowhead-shaped tissue profile and formation of sharp neuroectoderm tissue boundaries at the end of gastrulation (10 hpf) (Fig. 1b). Collectively, these observations indicate that the early ANP is dramatically reshaped during late gastrula.

**Figure 1.**
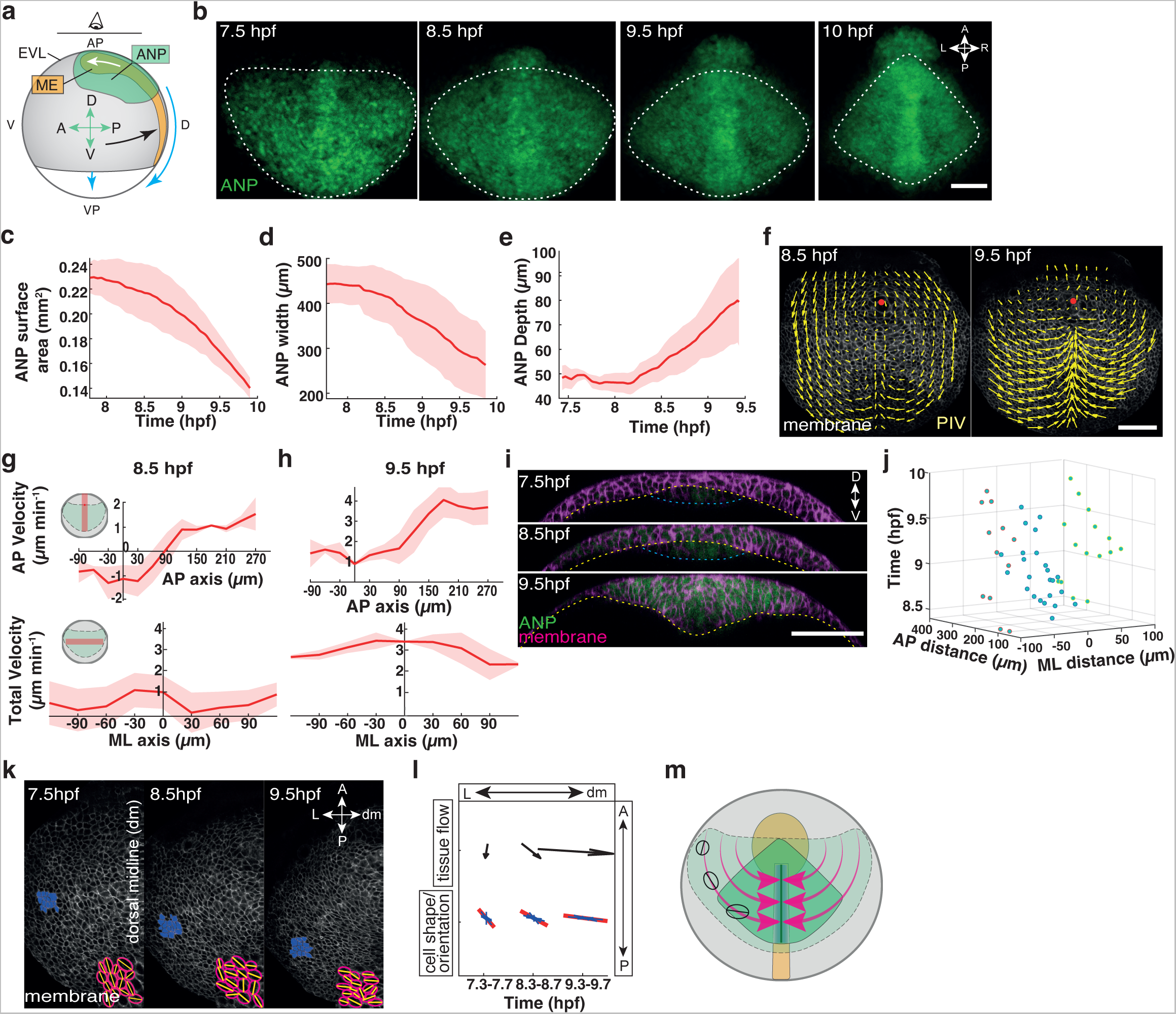
Anterior neural plate reshaping during gastrulation is driven by global tissue flows. **a)** Schematic illustrating positions and movements of anterior neural plate (ANP, green), mesendoderm (ME, orange) and epithelial enveloping layer (EVL) tissues in the zebrafish embryo during gastrulation. Lateral view, imaging angle (animal pole view) indicated. Animal–vegetal pole (AP–VP) and dorsal–ventral (D–V) indicate embryo orientation. Green arrows indicate ANP tissue orientation, anterior (A), posterior (P), dorsal (D) and ventral (V). White arrow, ME movement; black arrow, convergent extension (CE) movements; blue arrows, epiboly movements. **b)** Live imaging of neuroectoderm cells (maximum projection) in the ANP of *Tg(otx2:Venus)* wild type (wt) embryos during gastrulation (7.5–10 hours post fertilisation, hpf). White dotted line outlines the ANP. Anterior (A), posterior (P), left (L) and right (R) axes indicated. Scale bar, 100 µm. **c-e)** ANP tissue surface area (c), width (d) and tissue depth (e) against gastrulation time (hpf) in wt embryos. (n = 3 embryos). Mean ± s.e.m. **f)** Tissue flows of time-averaged velocities projected on membrane (mRFP) labelled neuroectoderm cells in the dorsal-most layer of the ANP in wt embryos at 8.5 hpf and 9.5 hpf (n = 3 embryos). Red dot indicates intersection of AP axis with anterior edge of ANP tissue. Scale bar, 100 µm. **g,h)** Cell velocities from (f) at 8.5 hpf (g) and 9.5 hpf (h) in anterior-posterior (AP) direction along the dorsal midline (position AP axis) and total velocities along the mediolateral (ML) axis (position ML axis) of the embryo (n = 3 embryos). AP velocity, X-axis: 0 marks anterior edge of the ANP; negative, positioned anterior and positive, positioned posterior. Y-axis: negative, posterior and positive, anterior-directed flows. Total velocity, X-axis: 0 marks the dorsal midline; negative, left and positive, right. Insets: illustration of location (red) of cells analysed in the embryo. Mean ± s.e.m. **i)** Confocal images (ventral view, transverse section) depicting the ANP in wt embryos through gastrulation. Cells membranes (mRFP, magenta) and ANP (otx2:Venus, green) labelled. Yellow dotted line demarcates the interface of the neuroectoderm and mesendoderm/yolk. Blue line indicates ventral edge of mesendoderm. Scale bar, 100 µm. **j)** Location and time of internalised cells within the ANP. Blue points represent cells along the dorsal midline. Red and green circled points represent cells on the right and left side of the embryo respectively. X-axis: distance left and right of the embryonic midline (0); Y-axis: distance posterior of the ANP leading edge; Z-axis: gastrulation stages (hpf). **k)** Confocal images of membrane-labelled (mRFP) neuroectoderm cells in the left half of the ANP during gastrulation (7.5/8.5/9.5 hpf). Orientation and shape of neuroectoderm cells in domains (highlighted in blue) originating from the lateral leading edge of the ANP at 7.5 hpf. Insets, cell outlines (red) as fitted ellipses and major cell axis (yellow line). Arrows indicate anterior-posterior (AP) and left-right (LR) axes. Scale bar, 100 µm. **l)** Orientation of tissue flow and cell orientation/shape within cell domains in (k). Arrows indicate the average direction and strengths of tissue flows over 20 minutes around the indicated times. Red lines, average cell orientation and shape (length) within this time interval. Blue lines, histogram of orientations of individual cells. (n = 3 embryos). **m)** Schematic of ANP tissue (green, animal pole view) shape changes and underlying mesendoderm (orange) during gastrulation. Arrows (magenta) highlight flows towards the dorsal midline and changes in cell orientation/shape are indicated at different time points.

### Local cell internalisation drives global tissue flows to reshape the ANP

To understand how cellular processes within the ANP contribute to the observed global tissue shape changes, we evaluated morphogenetic processes on a cellular level in cell membrane (mRFP) labelled *Tg(otx2:Venus)* zebrafish embryos to image neuroectoderm cell movements and shape changes (Supplementary Video 1). Quantitative analysis of cellular dynamics in curved 3D tissues, such as the neural plate, is an intricate problem and requires a robust methodology to transform 3D volumes into planar two-dimensional (2D) layers. To quantify local cellular processes across the entire ANP, we developed a computational image analysis workflow that enabled us to identify and align spatiotemporal landmarks in the embryo (ANP anterior edge and AP axis) and to compare cell and tissue dynamics across different imaging series and embryos (Supplementary Fig. 1c,d). This methodology also allowed us to visualise individual layers of the ANP tissue (superficial and deep cells) in the same plane by calculating planar projections of the whole 3D imaging volume (Supplementary Fig. 1e–i). This mapping process supported a downstream automated 2D image analysis of movements and segmentation of cells within the ANP (Supplementary Fig. 1h,i).

To quantify cell movements, we applied particle image velocimetry (PIV) to membrane labelled neuroectoderm cells in wild type (wt) embryos. Our analysis revealed the appearance of tissue flows with vortices on either side of the dorsal midline in the ANP at mid-gastrulation (8.5 hpf) (Fig. 1f), similar to flows previously identified in the early gastrula ^31^. We further identified the presence of opposing cell movements along the dorsal midline of the embryo with posterior directed (rearward) and anterior directed (forward) flows that collided at a stagnation point located within the ANP tissue (~90µm posterior of the ANP leading edge along the AP axis) (Fig. 1f,g). Rotational flows subsided after 8.75 hpf, at which time neuroectoderm cells displayed increasing mediolateral (ML) convergence movements towards the dorsal midline (Fig. 1f,h), which coincided with rapid narrowing of the tissue (Fig. 1b,d). Notably, lateral cells approaching the midline internalised (dorsal to ventral, DV), resembling tissue folding, and subsequently moved anteriorly (Fig. 1e,i). Internalisation was centred around the dorsal midline and occurred sequentially along the AP axis with the first appearance located posterior of the ANP leading edge (167 ± 9 µm s.e.m.; n = 3 embryos) (Fig. 1j). This was followed by a successive zipper-like anterior-to-posterior inward movement of neuroectoderm cells along the midline, suggesting that internalisation occurs stepwise rather than as a bulk movement (Fig. 1j; Supplementary Fig. 1j). Given that the timing of neuroectoderm cell internalisation coincided with prominent changes in flow topologies, we hypothesised that internalisation may generate forces affecting tissue flows. To address this, we measured changes in orientation and shape of neuroectoderm cells moving from the anterior-lateral edges of the tissue to the dorsal midline. We found that the local orientation of cells (direction of major axis) dynamically aligned with the direction of the regional flow after internalisation and also became progressively stretched over time (Fig. 1k,l; Supplementary Fig. 1k). Taken together, these observations indicate that internalisation along the DV axis may generate forces that control large scale tissue flows along the AP/ML axes, enabling gradual symmetrical tissue reshaping (Fig. 1m).

### Differential behaviours of mesendoderm cells initiate ANP tissue folding and cell internalisation

Our observations indicate that a folding event in the ANP leads to 3D reorganisation of the tissue and neuroectoderm cell internalisation. Tissue folding can be driven by local mechanical instabilities which can be instigated locally by regional cellular actomyosin contractility ^33–35^. Hence, we first investigated whether myosin 2 activity was enriched in cells at the dorsal side by staining for phosphorylated myosin 2 in neuroectoderm cells (8–9 hpf). Interestingly, we found no indication of increased areas of myosin 2 activity in neuroectoderm cells at or around the region of internalisation despite clear detection of elevated phosphorylated myosin 2 levels in underlying migrating axial mesendoderm cells (Supplementary Fig. 2a). We next investigated whether neuroectoderm cells undergo any shape changes during internalisation (Supplementary Video 2). We noticed that cells become progressively stretched along the direction of internalisation with increasing cell depth (Fig. 2a,b; Supplementary Fig. 2b). Interestingly, apical and basal surface areas of internalising cells were isotropically reduced during cell elongation (Supplementary Fig. 2c), suggesting that cell stretching is likely driven by ventral pulling forces or/and posterior pushing forces rather than cell intrinsic forces such as apical constriction.

**Figure 2.**
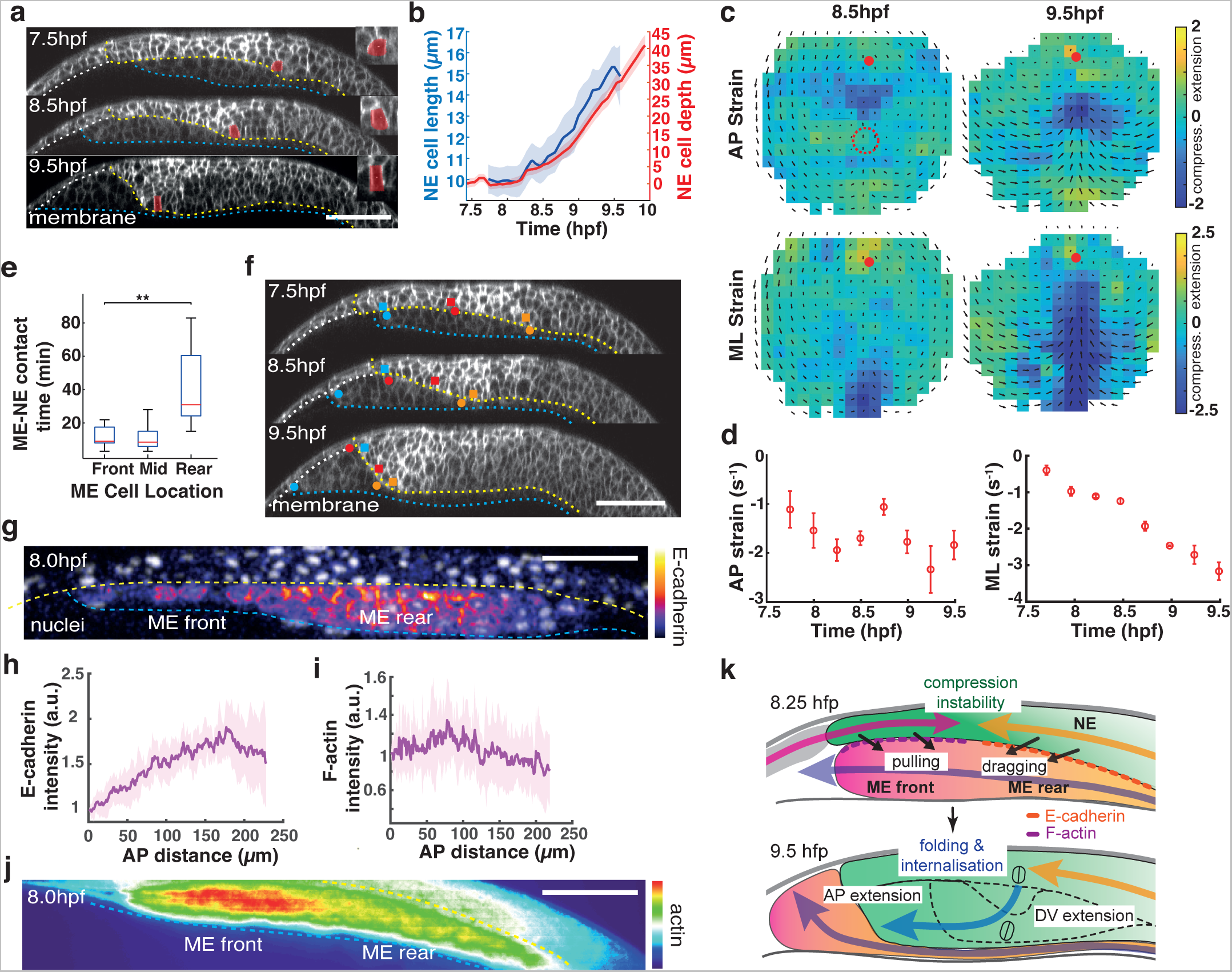
Differential behaviours of mesendoderm cells initiate anterior neural plate folding and internalisation. **a)** Confocal images (sagittal sections) representing the anterior neural plate (ANP) in wild type (wt) embryos through gastrulation (7.5/8.5/9.5 hours post fertilisation, hpf). Anterior (A) left, posterior (P) right, dorsal (D) up and ventral (V) down. Cell membranes (mRFP, white) labelled. Yellow line outlines neuroectoderm ventral and anterior border to mesendoderm, blue line indicates mesendoderm/yolk interface and white line marks non-neuroectodermal tissue. Representative internalising cell highlighted in red illustrate shape changes over time. Scale bar, 100 µm. **b)** Internalising neuroectoderm (NE) cell depth (red) and lateral cell length (blue) during gastrulation (n = 3 embryos). Mean ± s.e.m. **c)** Time-averaged ANP domain strain rates along the anterior-posterior (AP, upper panels) and mediolateral (ML, lower panels) axes of wild type embryos (n = 3 embryos). Average normal strain rate is colour-coded (minimum, green (0); maximum stretch, yellow; maximum compression, dark blue). Time-averaged tissue flows are indicated. Red dot marks intersection of ANP leading edge with AP axis. Dotted circle indicates initial position of internalisation. **d)** Maximum absolute ANP strain rates along the AP (left panel) and ML (right panel) axes of wt embryos as a function of time during gastrulation (plotted in 15 min intervals; n = 3 embryos). Negative values indicate compression. Mean ± s.e.m. **e)** Contact time between mesendoderm and neuroectoderm cells at different locations [marked in (f)] along the AP axis (n = 3 embryos). Kruskal–-Wallis test (**, p < 0.01). Mean ± s.e.m. **f)** Confocal images (sagittal sections) showing evolution of selected mesendoderm (circles) and neuroectoderm (squares) cells at the mesendoderm front (blue), mid (red) and rear (orange) during gastrulation. Orientation and labels as in (a). Scale bar, 100 µm. **g)** Live imaging (sagittal section, 8 hpf) of *Tg(gsc:cdh1-EGFP)* embryos showing nuclear label (white) and cadherin-1 (E-cadherin) expression levels (Fire LUT colour code; white maximum, black minimum) in the mesendoderm (ME). Orientation and labels as in (a). Scale bar, 100 µm. **h)** Distribution of E-cadherin in mesendoderm cells adjacent to neuroectoderm interface (g) as measured by intensity levels (a.u.) of gsc:cdh1-GFP signal along the front-rear (AP) axis (0=leading edge) (n = 3 embryos). Mean ± s.e.m. **i)** Distribution of F-actin in mesendoderm cells adjacent to neuroectoderm interface (j) as measured by intensity levels (a.u.) of lifeact:EGFP signal along the front-rear (AP) axis (0=leading edge) (n = 3 embryos). Mean ± s.e.m. **j)** Live imaging (maximum projection sagittal sections, 8 hpf) of *Tg(actb1:lifeact-EGFP)* embryos showing F-actin expression levels (Thermal LUT colour code; red maximum, dark blue minimum) in the mesendoderm. Orientation and labels as in (a). Scale bar, 100 µm. **k)** Schematic (lateral view) illustrating forces (black arrows) exerted by the migrating mesendoderm (ME, pink/orange) leading to opposing movements (pink arrow, posterior and orange arrow, anterior-directed) in the ANP (green), generating instabilities and folding enabling neuroectoderm (NE) cell internalisation and tissue extension along the dorsal-ventral (DV) and anterior-posterior (AP) axes. E-cadherin (orange dashed line), F-actin (purple dashed line) and cell shapes depicted.

To analyse such external forces that might locally trigger mechanical instabilities and hence tissue folding spatiotemporally, we then mapped local tissue strain rates as readout of force-driven deformations ^36^. Notably, we identified an area of localised, progressively increasing compression at the dorsal midline occurring before internalisation (7.5–8.25 hpf) (Fig. 2c,d; Supplementary Fig. 2d). This corresponded with the flow stagnation point along the AP axis (Fig. 1e,f), suggesting that opposing cell movements locally compact the tissue. We further observed that neuroectoderm cell internalisation initiated at a point posterior to the compressed area (Fig. 2c), suggesting that local tissue compaction likely acts as a hinge point around which the tissue can pivot, enabling inward folding (Supplementary Video 2). Furthermore, our mediolateral (ML) strain analysis revealed that internalisation was accompanied by progressively increasing tissue compression on either side of the dorsal fold region (9.5 hpf; Fig. 2c,d), suggesting that internalisation along the DV axis may produce significant forces accounting for tissue convergence. Taken together, these measurements suggest that local deformations in the ANP likely result from multiple stresses acting in a spatiotemporally controlled manner.

We next focused on understanding how stresses leading to opposing cell movements in the ANP might be generated to initiate folding. To address this, we first analysed the behaviour of underlying axial mesendoderm progenitor (prechordal plate) cells along the tissue interface to determine the duration of adhesion between mesendoderm and overlying neuroectoderm cells (Supplementary Video 2). Strikingly, we found that mesendoderm cells located at the front of the collective showed frequent contact exchanges with neuroectoderm cells, reminiscent of stick-slip motion typically observed in migrating cells on a substrate (Fig. 2e,f). In contrast, mesendoderm cells positioned at the rear of the collective and in the posterior positioned notochord displayed long-lasting adhesions to overlying neuroectoderm cells (Fig. 2e,f). We next explored whether variations in E-cadherin organisation in the mesendoderm collective would explain such behaviours ^31, 37^. Thus, we generated a *Tg(gsc:cdh1-EGFP)* transgenic line which allowed live imaging of E-cadherin (cadherin 1) dynamics specifically in the axial mesendoderm of embryos (Fig. 2g). Interestingly, we observed a greater accumulation of E-cadherin in rear than front mesendoderm cells (Fig. 2g,h) which was independent of promoter-driven expression levels (Supplementary Fig. 2e). To explore differences in migratory behaviour, we imaged F-actin processes in mesendoderm cells and found that mesendoderm cells positioned at the front of the collective showed substantial F-actin activity, often enriched at protrusions at the neuroectoderm interface, whereas cells at the rear displayed reduced F-actin levels (Fig. 2i,j; Supplementary Fig. 2f). To further substantiate whether mesendoderm front cells might be responsible for posterior directed movements of neuroectoderm cells, we measured neuroectoderm cell displacements around the animal pole where we expected minimal epiboly movements^38^ and hence any substantial movements in the neuroectoderm to be driven by adjacent mesendoderm cells. In support of this hypothesis, we found that neuroectoderm cells at the animal pole moved towards the approaching mesendoderm in a distance dependent manner (Supplementary Fig. 2g), suggesting that posterior directed neuroectoderm movements likely originate from the underlying mesendoderm. Together, these findings indicate that opposing movements in the ANP likely arise from differential migration and adhesion behaviours in the mesendododerm, whereby mesendoderm front cells exert pulling (traction) forces and rear cells drag neurectoderm anterior, in agreement with recently reported frictional forces acting between the tissues^31^. We propose that an interplay of these forces is sufficient to account for local mechanical instabilities in the ANP to initiate tissue folding (Fig. 2k).

### A spatiotemporal interplay of forces predicts global tissue flows in the ANP

Our observation that global flow patterns change during gastrulation suggests that neuroectoderm tissue flows are regulated by multiple forces that vary in space and time. To address this, we developed a mechanical model of neuroectoderm tissue flows to determine the minimal set of local forces required to reproduce tissue flows during different gastrulation stages (8.25–10 hpf) (Fig. 3a–j). On symmetrising (i.e., left-right averaging) experimental flow data, we found that flows before internalisation (8.25 hpf) exhibited vortices on either side of the dorsal midline and a nearby extension point on the midline (Fig. 3a). After the initiation of internalisation (8.75 hpf), the symmetrised flows changed and displayed a turning flow on either side of the midline and a sink on the midline that is the signature of internalised cells leaving the plane of the ANP (Fig. 3f). To reproduce these flow topologies *qualitatively*, we modelled the neuroectoderm as an infinite two-dimensional incompressible viscous fluid, in which flows are driven by point force and point sink singularities ^39^ on an axis representing the dorsal midline (Supplemental Note).

**Figure 3.**
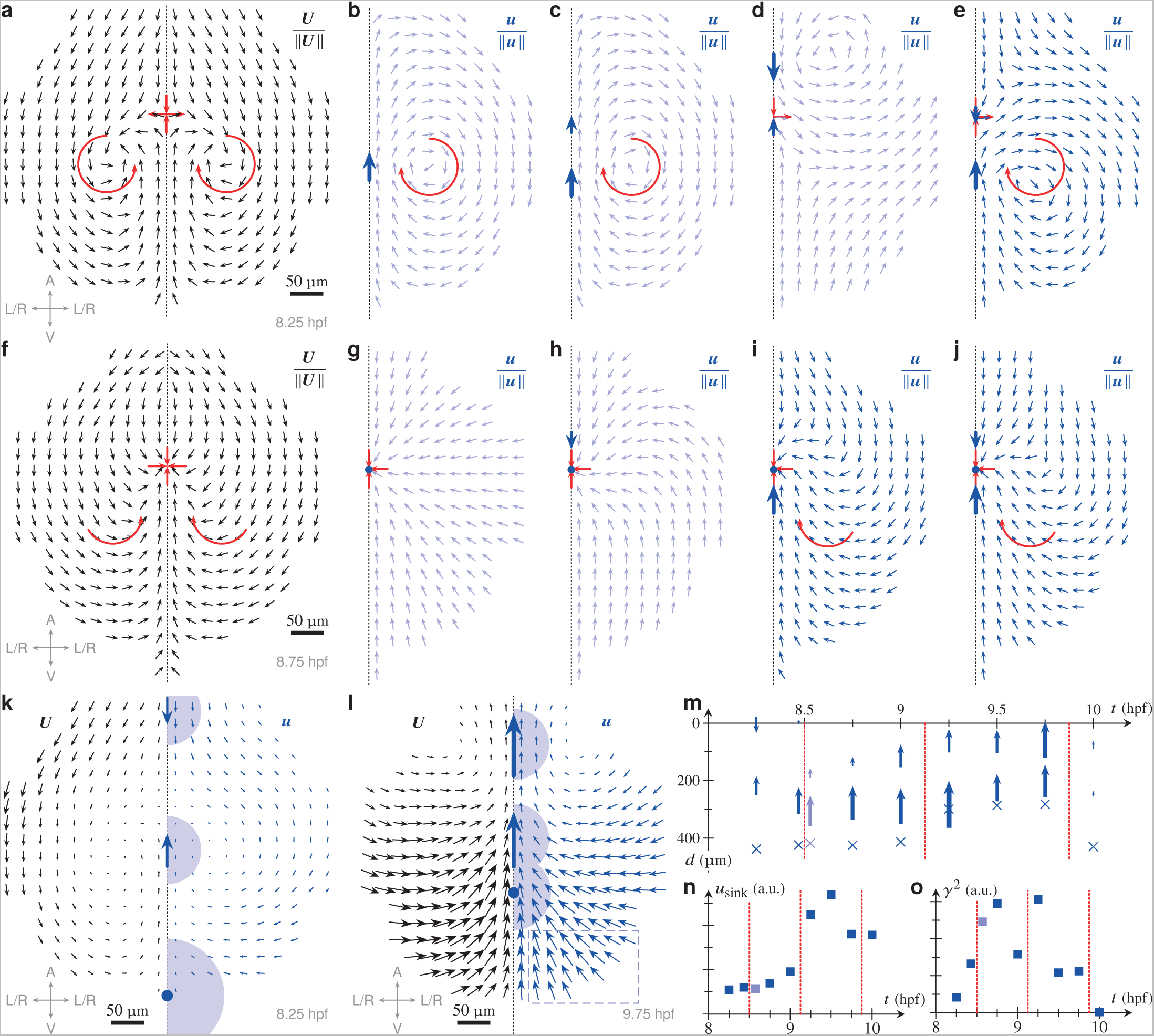
A mechanical model predicts a spatiotemporal interplay of forces controlling tissue flows in the anterior neural plate. **a)** Symmetrised experimental flow directions *U*/‖*U*‖ before internalisation (8.25 hours post fertilisation, hpf); the main features of the flow are lateral vortices and a nearby extension point along the dorsal midline (highlighted red). Inset: embryo axes (A: animal, V: vegetal, L/R: left-right symmetrised). Scale bar, 50 µm. **b)** Flow directions ***u***/‖***u***‖ resulting from a single force in the anterior direction. The flow features a lateral vortex but does not have an extension point nearby. **c)** Flow directions *u*/‖*u*‖ resulting from two parallel force singularities; the flow still features a lateral vortex but no extension point. **d-e)** Flow directions *u*/‖*u*‖ resulting from two antiparallel force singularities. (d) If the posterior-directed force dominates, the flow displays an extension point but incorrect vortices. (e) If the anterior-directed force is larger, extension point and vortices are correct. **f)** Symmetrised experimental flow directions *U*/‖*U*‖ after internalisation has started (8.75 hpf). The main features of the flow (which has lost the vortices) are a sink on the axis and a turning flow posterior to the sink. Inset: embryo axes as in (a). Scale bar, 50 µm. **g)** Flow directions *u*/‖*u*‖ resulting from a single sink singularity lack the turning flow. **h)** Flow directions *u*/‖*u*‖ resulting from the combination of a sink and a pulling force; the orientation of the turning flow is incorrect. **i)** Flow directions *u*/‖*u*‖ resulting from the combination of a sink and a drag force feature a sink and a turning flow of the correct orientation. **j)** Flow directions *u*/‖*u*‖ resulting from a combination of a sink, a drag force and a smaller pulling force feature a sink and a correct turning flow, too. **k)** Experimental flow field *U* (left) and fitted flow field *u* (right) at 8.5 hpf. Arrows indicate force directions and positions; the dot indicates the sink position. Shaded areas indicate the characteristic areas over which the singularities are smeared out. The fitted sinks represent contributions from both compressibility and internalisation. Inset: embryo axes as in (a). Scale bar, 50 µm. **l)** Experimental flow field *U* (left) and fitted flow field *u* (right) at 9.75 hpf, similar to (k) Rectangle emphasises that the fit underestimates the left-right flows in the posterior region. Inset: embryo axes as in (a). Scale bar, 50 µm. **m)** Plot of the positions of the regularised force singularities (arrows) and regularised sinks (dots) against time; positions are given in terms of the distance *d* with respect to the anterior limit of the tissue, as determined by the experimental data. Sizes of arrows indicate force magnitudes. Different fits of very similar fit scores shown for 8.5 hpf. Vertical red lines separate different mechanical regimes. **n)** Plot of the fitted sink velocity *u*_sink_ (normalised with its mean value) against time. Vertical red lines show mechanical transitions, as in (m). A.u.: arbitrary units. **o)** Plot of the fitted friction coefficient *γ*^2^ (normalised with its mean value) against time. Vertical red lines show mechanical transitions, as in (m). A.u.: arbitrary units.

First, we asked whether forces acting on the neurectoderm could reproduce the flow topology observed *in vivo* before internalisation (8.25 hpf) (Fig. 3b–e). We found that neither a single force singularity, nor two parallel ones could account for the observed topological features (Fig. 3b,c). However, two opposing force singularities acting on the neurectoderm along the dorsal midline were able to account for the observed tissue flows before internalisation (Fig. 3d,e). Strikingly, this model predicts that the anterior-directed force must be stronger than the posterior-directed one to reproduce the observed orientation of the vortices (Fig. 3e). Next, we modelled the flows after initiation of tissue folding (8.75 hpf) (Fig. 3g–j). To represent forces directed out of the ANP tissue plane that led to internalisation of cells at the midline, we introduced a single sink singularity on the AP axis. We found that a sink on its own (Fig. 3g), or in combination with a force directed towards the posterior (Fig. 3h) could not reproduce the experimental flow topology (Fig. 3f). However, combinations of a sink and a single force directed towards the anterior (Fig. 3i) or two opposing forces (Fig. 3j) reproduced this topology.

Our qualitative model thus indicates two mechanical regimes: flows before tissue internalisation result from opposing forces along the AP tissue axis, while an anterior-directed force and an internalisation force out of the tissue plane drive flows during internalisation. To test this hypothesis, we sought to show that the minimal combination of two forces and one sink that we have shown above to be *necessary* to reproduce the flow topologies *qualitatively*, is in fact sufficient to reproduce the experimentally measured flows *quantitatively*. For this purpose, we extended our model (Supplemental Note) by including friction between the neuroectoderm and the overlying enveloping layer (EVL) to remove the Stokes paradox ^39^ and regularising the divergences of the force and sink singularities, smearing them out over finite distances that represent the characteristic physical extents over which they are applied to the tissue. We thus fitted the positions, magnitudes, directions, and extents of two such regularised forces and one such regularised sink and the EVL-neurectoderm friction to the experimentally quantified tissue flows for different timepoints (8.25–10 hpf) (Fig. 3k–o and Supplemental Note).

Importantly, this minimal mechanical model could indeed quantitatively reproduce the experimentally observed neuroectoderm tissue flows (Fig. 3k,l). The fits revealed multiple mechanical transitions during gastrulation (8.25–10 hpf) (Fig. 3m–o) and in particular differences in the directions, relative positions, and magnitudes of the forces and sink driving the flows (Fig. 3m–o). Fits for early time points before internalisation (< 8.5 hpf) featured two opposing forces along the AP axis (the smaller posterior-directed, the larger anterior-directed) (Fig. 3k,m), agreeing with predictions from our qualitative model (Fig. 3a–e). The model predicted a first mechanical transition when internalisation begins (~8.5 hpf) and the two opposing forces gave way to two parallel forces directed towards the anterior (Fig. 3m), with a simultaneous increase in EVL-neuroectoderm friction (Fig. 3o). Consistently with the start of internalisation, the fitted sink velocity increased (Fig. 3n) and reached a maximum during a second mechanical transition during late gastrulation (~9 hpf) (Fig. 3n). Interestingly, forces repositioned towards the leading edge of the tissue at that time (Fig. 3m), indicating positional dynamics of the exerted forces. During late gastrulation (~10 hpf), the fitted sink strengths decreased, and the forces and friction were considerably reduced (Fig. 3m,o), suggesting a final mechanical relaxation as the ANP acquires its final shape. In summary, we have shown that the minimal mechanics identified by our qualitative model (Fig. 3a–j) are sufficient to reproduce the experimental flows quantitatively (Fig. 3k,l). Moreover, our quantitative model revealed how mechanical transitions at different developmental stages lead to a spatiotemporal force distribution regulating the experimentally measured flows.

### Mesendoderm migration is essential for internalisation and tissue folding

Based on our *in vivo* and *in silico* analyses, we hypothesised that the underlying mesendoderm may act as a key force generator driving ANP shape changes. To verify this, we investigated ANP morphogenesis in the absence of mesendoderm cells and imaged neuroectoderm cell dynamics in *lefty1* mRNA injected *Tg(oxt2:Venus)* zebrafish embryos (Supplementary Video 3). We observed that, in these embryos, the ANP was misshaped and remained close to the embryonic margin (Fig. 4a; Supplementary Fig. 3a), consistent with previous observations of stained tissue in MZ*oep* mutant embryos ^31, 40^. In contrast to wt embryos, we observed that the tissue width was nearly unaltered in *lefty1* embryos during gastrulation (Fig. 4b; Supplementary Fig. 3b). Importantly, we detected no characteristic internalisation of neuroectoderm cells (Fig. 4c,d; Supplementary Fig. 3c) or rotational flow patterns (Fig. 4e) in embryos lacking mesendoderm. Further, *lefty1* embryos showed wide-ranging ANP tissue compression along the AP axis without significant ML deformations, driven by unidirectional posterior flows of neuroectoderm cells against the margin (Fig. 4f,g; Supplementary Fig. 3d,e). Consequently, neuroectoderm cells became oriented perpendicular to the tissue flow along the AP tissue axis (Supplementary Fig. 3f). Together, this indicated that axial mesendoderm migration provides essential extrinsic forces to regulate internalisation and tissue flows to reshape the ANP during gastrulation.

**Figure 4.**
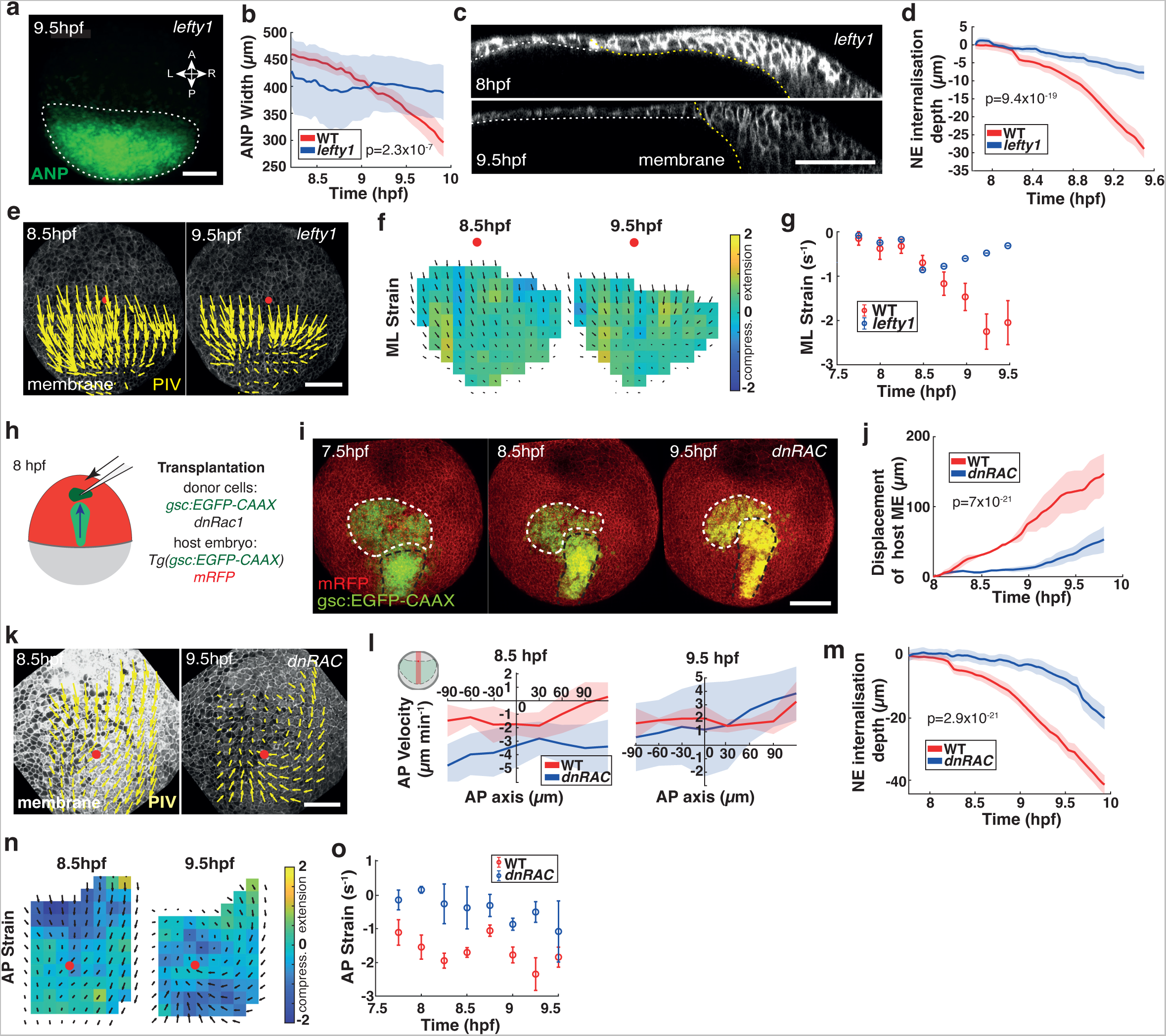
Mesendoderm migration is essential for anterior neural plate folding and internalisation. **a)** Live imaging of neuroectoderm cells (maximum projection) in the anterior neural plate (ANP) of *Tg(otx2:Venus) lefty1* morphant embryos at 9.5 hours post fertilisation (hpf). White dotted line outlines the ANP. Scale bar, 100 µm. **b)** ANP tissue width of neuroectoderm cells in wt and *lefty1* morphant embryos over time of gastrulation. (n = 3 embryos). Mean ± s.e.m. **c)** Confocal images (sagittal sections) depicting the ANP in *lefty1* morphant embryos at 8 hpf and 9.5 hpf. Dorsal up, ventral down, anterior to the left and posterior to the right. Cell membranes (mRFP, white) labelled. Yellow line marks interface between neuroectoderm and yolk, white line outlines non-neuroectodermal tissue. Scale bar 100 µm. **d)** Internalisation depth of neuroectoderm cells in wt and *lefty1* morphant embryos over time of gastrulation. (n = 3 embryos). Mean ± s.e.m. **e)** Tissue flows of time-averaged velocities projected on membrane (mRFP) labelled neuroectoderm cells in the dorsal-most layer of the ANP in *lefty1* morphant at 8.5 hpf and 9.5 hpf (n = 3 embryos). Red dot indicates intersection of AP axis with anterior edge of ANP tissue. Scale bar, 100 µm. **f)** Time-averaged ANP domain strain rates along the mediolateral (ML) axis of *lefty1* morphant embryos (n = 3 embryos). Average normal strain rate is colour-coded (minimum, green (0); maximum stretch, yellow; maximum compression, dark blue). Time-averaged tissue flows are indicated. Red dot marks intersection of ANP leading edge with AP axis. **g)** Maximum absolute strain rates along the ML axis of wt and *lefty1* embryos as a function of time during gastrulation (plotted in 15 min intervals; n = 3 embryos). Negative values indicate compression. Mean ± s.e.m. **h)** Schematic illustrating transplantation assay. Donor dnRac1 expressing mesendoderm cells (dark green) transplanted anterior to host mesendoderm (light green) before internalisation at 8 hpf. **i)** Live imaging (animal pole view, ventral side up, dorsal side down) of *Tg(gsc:GFP-CAAX)* host embryo transplanted with dnRac1 (*dnRAC*) expressing mesendoderm donor cells (*gsc:GFP-CAAX*). Donor mesendoderm (white dashed outline) was transplanted anterior to host mesendoderm (black dashed outline) at 7.5 hpf and imaged throughout gastrulation. Membrane (mRFP) labelled in red. Scale bar, 100 µm. **j)** Anterior displacement of mesendoderm in wt (red) and transplanted (*dnRAC*, blue) embryos (n = 3 embryos). Error bar, s.e.m. **k)** Tissue flows of time-averaged velocities projected on membrane (mRFP) labelled neuroectoderm cells in the dorsal-most layer of the ANP in transplanted embryos at 8.5 hpf and 9.5 hpf (n = 3 embryos). Red dot indicates intersection of AP axis with anterior edge of ANP tissue. Scale bar, 100 µm. **l)** Cell velocities from (k) at 8.5 hpf and 9.5 hpf in anterior-posterior (AP) direction along the dorsal midline of wt (red) and transplanted (*dnRAC*, blue) embryos (n = 3 embryos). AP velocity plots, X-axis (µm): 0 marks anterior edge of the ANP; negative, anterior and positive posterior positioned. Y-axis (µm/min): negative, posterior and positive, anterior-directed flows. Mean ± s.e.m. **m)** Neuroectoderm internalisation depth (µm) of dorsal layer of neuroectoderm cells in wt (red) and transplanted (*dnRAC*, blue) embryos (n = 3 embryos). Mean ± s.e.m. **n)** Time-averaged ANP domain strain rates along the AP axis of wt and *dnRAC* morphant embryos (n = 3 embryos). Average normal strain rate is colour-coded (minimum, green (0); maximum stretch, yellow; maximum compression, dark blue). Time-averaged tissue flows are indicated. Red dot marks intersection of ANP leading edge with AP axis. **o)** Maximum absolute ANP strain rates (mean) along the AP axis of wt embryos as a function of time during gastrulation (plotted in 15 min intervals; n = 3 embryos). Negative values indicate compression. Mean ± s.e.m.

We further reasoned that blocking mesendoderm migration in wt embryos may eliminate the anterior drag of neuroectoderm cells and affect the generation of instabilities required for tissue folding. To address this, we inhibited the small Rho GTPase Rac1 in mesendoderm cells, an essential protein for protrusion formation in mesendoderm cells ^41^. We locally transplanted mesendoderm cells expressing dominant negative Rac1 (dnRac) from donor embryos to the leading edge of mesendoderm in wt host embryos before neuroectoderm internalisation occurred (7.5–8 hpf) (Fig. 4h,i). Notably, when endogenous mesendoderm cells joined the migration-deficient transplanted cells, they stalled in their anterior migration (Fig. 4j; Supplementary Fig. 3g). Consequently, the neuroectoderm only displayed unidirectional posterior-directed flows along the dorsal midline at 8.5 hpf, suggesting that forces controlling anterior-directed neuroectoderm movements were abolished (Fig. 4k,l; Supplementary Fig. 3j). Notably, we did not observe any tissue folding and internalisation under these conditions (Fig. 4m; Supplementary Fig. 3h,i). Strain maps revealed that stresses in the ANP were altered and local AP tissue compaction at 8.5 hpf as well as lateral compression due to internalisation at 9.5 hpf were absent (Fig. 4n,o; Supplementary Fig. 3k,l). However, we noticed that internalisation movements were partially restored by late gastrulation (9.5 hpf) (Fig. 4m; Supplementary Fig. 3i) when the mesendoderm regained momentum (Fig. 4j), likely through AP axis extension of posterior tissues. Overall, we conclude that mesendoderm migration during gastrulation constitutes a major force generating mechanism providing extrinsic forces critical for tissue folding and correct timing of the initiation of cell internalisation.

### Convergent extension movements are dispensable for internalisation but required for ANP tissue extension

Our model prediction revealed a second anterior directed force acting at the posterior end of the ANP during late gastrulation (Fig. 3l). We hypothesised that this force may be associated with convergent extension (CE) which are highly conserved global cell movements essential for embryonic axis formation and generation of tissue-scale force production ^42, 43^. To address how CE contributes to ANP tissue morphogenesis, we generated *Tg(otx2:Venus); wnt11* (*silberblick*)^44^ transgenic mutants to image ANP formation in embryos lacking CE movements (Supplementary Video 4). Measurements of live tissue dynamics in these embryos revealed that the ANP is only partially reshaped during gastrulation and remains laterally expanded compared to wt embryos (Fig. 5a,b; Supplementary Fig. 4a,b). We next analysed cellular rearrangements within the neuroectoderm to identify whether changes in internalisation and/or tissue flows could account for defective ANP shape. Strikingly, we found that the timing and spatial location of initiation of internalisation are largely preserved in *wnt11* mutant embryos (distance posterior of ANP leading edge: *wnt11*: 181 ± 12 µm s.e.m; wt: 167 ± 9 µm s.e.m.; n = 3 embryos each) (Fig. 5c), regardless of the slight decrease in mesendoderm velocity (Supplementary Fig. 4c,d). However, internalisation in *wnt11* mutants showed an increased neuroectoderm internalisation depth (Fig. 5d; Supplementary Fig. 4e). Furthermore, the range of anterior displacement of the internalised tissue was drastically reduced and remained farther posterior throughout gastrulation compared to wt embryos (Fig. 5e), suggesting that CE supports AP extension of the ventral ANP tissue. To test whether this limited range of internalised tissue motion would affect force generation, we analysed neuroectoderm tissue flows in *wnt11* mutant embryos. We found that flow topologies and cell orientations along the flow in *wnt11* mutants were qualitatively similar to the wt at 8.5 hpf (Fig. 5f; Supplementary Fig. 4f,g). However, flow velocities along the AP, and especially the ML axis, were considerably reduced in stages following internalisation (9.5 hpf; Fig. 5f,g). Remarkably, we found that tissue deformations and strain rates in *wnt11* mutant embryos were comparable to the wt (Fig. 5h,i; Supplementary Fig. 4h,i). Yet, neuroectoderm cell internalisation remained locally restricted to the posterior of the ANP in *wnt11* mutants (Fig. 5c) and lacked the typical internalisation pattern along the AP axis observed in wt embryos (Supplementary Fig. 1j). Interestingly, *wnt11* mutant embryos displayed a kink in the ANP on the dorsal site of internalisation (Supplementary Fig. 4j), indicating that ventral tissue extension supported by CE is integral to preserve proper ANP tissue geometry. Collectively, we conclude that CE movements are dispensable for internalisation but essential to maintain robust tissue flows and ANP reshaping by mechanically supporting ventral tissue extension following internalisation.

**Figure 5.**
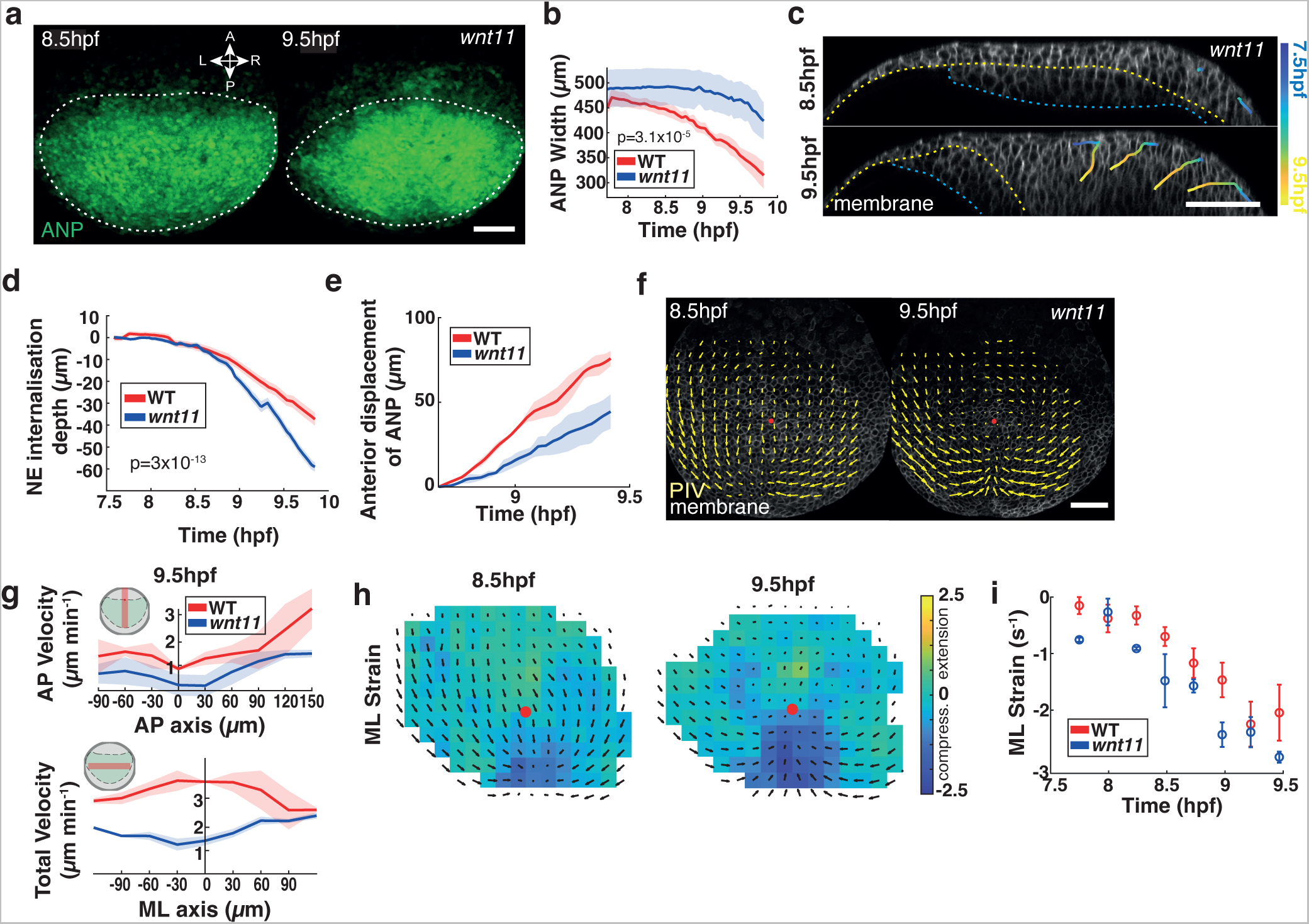
Convergent extension is integral for anterior neural plate shape by supporting tissue extension. **a)** Live imaging of neuroectoderm cells (maximum projections) in the anterior neural plate (ANP) of *Tg(otx2:Venus); wnt11* mutant embryos during gastrulation (8.5–9.5 hours post fertilisation, hpf). White dotted line outlines the ANP. Anterior (A), posterior (P), left (L) and right (R). Scale bar, 100 µm. **b)** ANP tissue surface width over time of gastrulation (hpf) in *wnt11* mutant embryos. (n = 3 embryos). Mean ± s.e.m. **c)** Confocal images (sagittal sections) showing representative cells tracked within the ANP in *wnt11* mutant embryos through gastrulation. Dorsal up, ventral down, anterior left and posterior right. Cell membranes (mRFP, white) labelled. Yellow dashed line: neuroectoderm/mesendoderm border, blue dashed line: mesendoderm/yolk interface. Colour code: 7.5 hpf (blue) to 9.5 hpf (yellow). Scale bar, 100 µm. **d)** Internalisation depth of neuroectoderm (NE) cells in wt and *wnt11* mutant embryos over time of gastrulation. (n = 3 embryos). Mean ± s.e.m. **e)** Anterior displacement of internalising neuroectoderm (NE) cells in wt (red) and *wnt11*(blue) mutant embryos. (n = 3 embryos) Mean ± s.e.m. **f)** Tissue flows of time-averaged velocities projected on membrane (mRFP) labelled neuroectoderm cells in the dorsal most layer of the ANP in *wnt11* mutant embryos at 8.5 hpf and 9.5 hpf (n = 3 embryos). Red dot indicates intersection of AP axis with anterior edge of ANP tissue. Scale bar, 100 µm. **g)** Cell velocities at 9.5 hpf in anterior-posterior (AP) direction along the dorsal midline and total velocities along the mediolateral (ML) axis of wt (red) and *wnt11* mutant (blue) embryos (n = 3 embryos). AP velocity plots, X-axis (µm): 0 marks anterior edge of the ANP; negative, anterior and positive posterior positioned. Y-axis (µm/min): negative, posterior and positive, anterior-directed flows. Total velocity plots, X-axis (µm): 0 marks the dorsal midline of the embryo; negative, left and positive right positioned (mediolateral, ML). Y-axis (µm/min): velocity. Mean ± s.e.m. **h)** Time-averaged ANP domain strain rates along the ML axis of *wnt11* mutant embryos (n = 3 embryos). Average normal strain rate is colour-coded (minimum, green (0); maximum stretch, yellow; maximum compression, dark blue). Time-averaged tissue flows are indicated. Red dot marks intersection of ANP leading edge with AP axis. **i)** Maximum absolute strain rates along the ML axis of wt and *wnt11* embryos as a function of time during gastrulation (plotted in 15 min intervals; n = 3 embryos). Negative values represent compression. Mean ± s.e.m.

### Internalisation depends on E-cadherin-mediated tissue interaction

Finally, we sought to identify how mechanical coupling between the neuroectoderm and mesendoderm is established to enable cell internalisation. Recent reports highlighted important roles for E-cadherin (cadherin-1) and N-cadherin (cadherin-2) in heterotypic tissue interactions between the zebrafish neural plate and the underlying mesendoderm ^31, 45^. To test for a functional role of both of these adhesion receptors in regulating internalisation of the neuroectoderm, we knocked-down the levels of both adhesion molecules individually. We found that ANP shape was not altered during gastrulation in N-cadherin depleted embryos (*cdh2* morphants) (Supplementary Fig. 5a–c). Further, *cdh2* morphants displayed similar neuroectoderm internalisation behaviour to wt (Supplementary Fig. 5d,e), indicating that N-cadherin is dispensable for shaping the ANP tissue during gastrulation.

We next moderately reduced the amount of E-cadherin in embryos (*cdh1* morphants) to avoid early deep cell epiboly arrest, and to allow for sufficient development until the end of gastrulation ^46, 47^. Live imaging of *cdh1* morphants (Supplementary Video 5) revealed very little change in ANP surface area or width suggesting strongly diminished ANP tissue remodelling (Fig. 6a–c; Supplementary Fig. 5f,g). Remarkably, neuroectoderm cells did not internalise and remained in superficial layers in these embryos (Fig. 6d,e; Supplementary Fig. 5h) with a strongly reduced tendency to elongate in the DV direction as observed in wt embryos (Fig. 6f; Supplementary Fig. 5i). We hypothesised that changes in inter-tissue force coupling should primarily affect mesendoderm cells at the rear responsible for dragging neuroectoderm cells anterior and measured the duration of heterotypic tissue adhesion. Indeed, we found that especially rear mesendoderm adhesion to neuroectoderm was significantly reduced in *cdh1* morphants (Fig. 6g). While opposing movements along the dorsal midline were still detectable in *cdh1* morphants at 8.5 hpf, velocities were substantially slower along the AP and ML axis of the tissue (Fig. 6h,i; Supplementary Fig. 5j). Further, local tissue compaction along the AP axis required for initiation of ANP folding at 8.5 hpf was absent (Supplementary Fig. 5k,l), as were ML tissue deformations along the dorsal midline were completely missing at 9.5 hpf (Fig. 6j,k). Taken together, our results suggest that E-cadherin, but not N-cadherin, mediated adhesion is critical to enable neuroectoderm internalisation by supporting inter-tissue force coupling.

**Figure 6.**
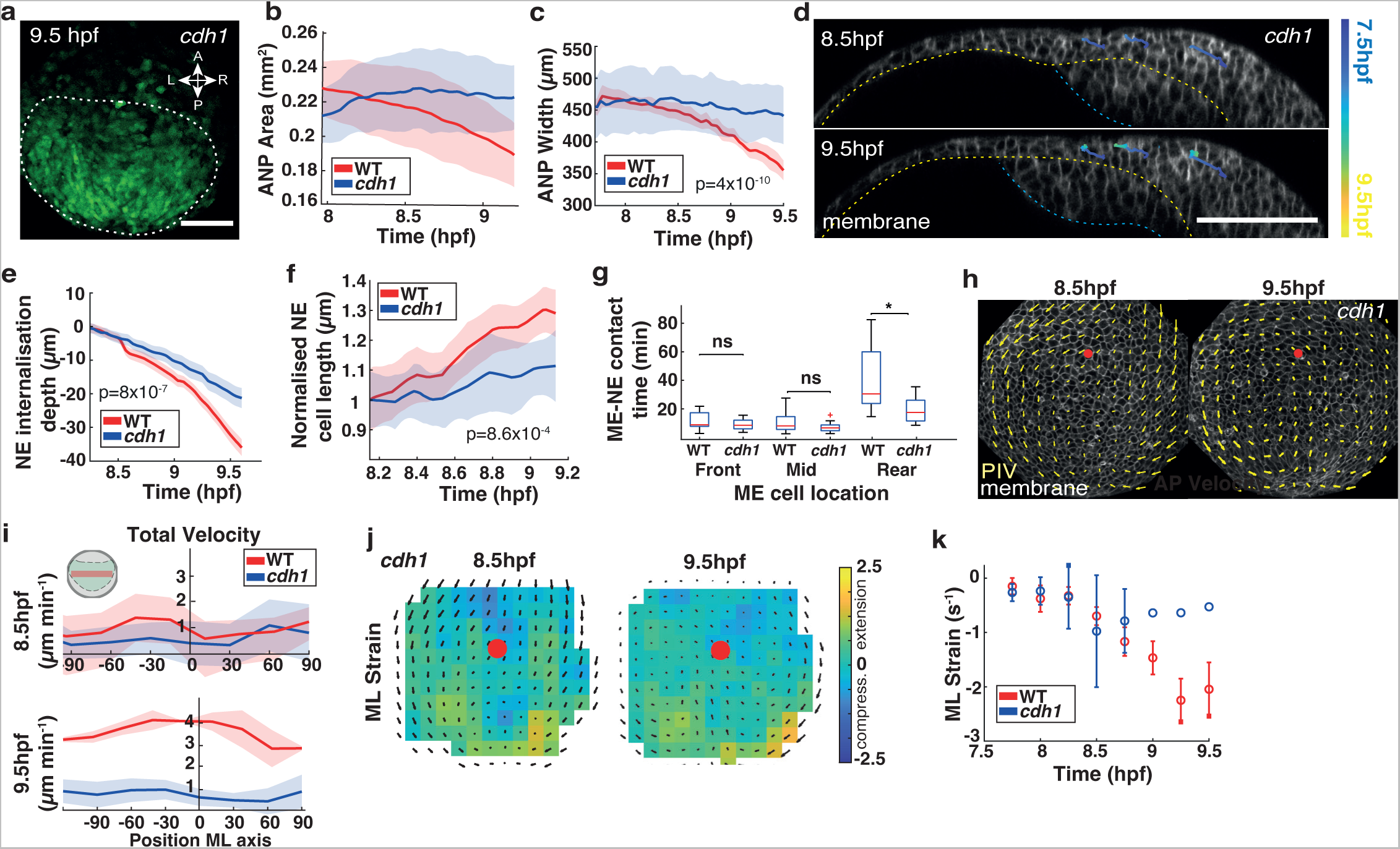
E-cadherin-mediated inter-tissue adhesion is required for neuroectoderm internalisation. **a)** Live imaging of neuroectoderm cells (maximum projections) in the anterior neural plate (ANP) of *Tg(otx2:Venus) cadherin-1 (cdh1)* morphant embryos at the end of gastrulation (9.5 hours post fertilisation, hpf). White dotted line outlines the ANP. Anterior (A), posterior (P), left (L) and right (R). Scale bar, 100 µm. **b,c)** ANP tissue surface tissue area **(b)** and tissue width **(c)** of wt (red) and *cdh1* morphant (blue) embryos during gastrulation. Mean ± s.e.m. **d)** Confocal images (sagittal sections) showing representative cells tracked within the ANP in *cdh1* morphant embryos through gastrulation. Dorsal up, ventral down, anterior left and posterior right. Cell membranes (mRFP, white) labelled. Yellow dashed line: neuroectoderm/mesendoderm border, blue dashed line: mesendoderm/yolk interface. Colour code: cells tracked from 7.5 hpf (blue) to 9.5 hpf (yellow). Scale bar, 100 µm. **e)** Neuroectoderm (NE) internalisation depth in wildtype (wt, red) and *cdh1* morphant (blue) embryos over time of gastrulation (hpf) (n = 3 embryos). **f)** Length of internalising NE cells in wt (red) and *cdh1* (blue) morphant embryos during gastrulation (n = 3 embryos). Mean ± s.e.m. **g)** Contact duration between and neuroectoderm (NE) and mesendoderm (ME) cells at different locations along the AP axis in *wt* and *cdh1* embryos (n = 3 embryos). Mean ± s.e.m. **h)** Tissue flows of time-averaged velocities projected on membrane (mRFP) labelled neuroectoderm cells in the dorsal most layer of the ANP in *cdh1* morphant embryos at 8.5 hpf and 9.5 hpf (n=3 embryos). Red dot indicates intersection of AP axis with anterior edge of ANP tissue. Scale bar, 100 µm. **i)** Total cell velocities (Y-axis, µm/min) from (h) along the mediolateral (ML) axis of wt (red) and *cdh1* (blue) embryos at 8.5 hpf and 9.5 hpf (n = 3 embryos). X-axis (µm): 0 marks dorsal midline; negative, left and positive, right positioned. Mean ± s.e.m. **j)** Time-averaged ANP domain strain rates along the mediolateral (ML) axes of *cdh1* morphant embryos (n = 3 embryos). Average normal strain rate is colour-coded (minimum, green (0); maximum stretch, yellow; maximum compression, dark blue). Time-averaged tissue flows are indicated. Red dot marks intersection of ANP leading edge with AP axis. **k)** Maximum absolute strain rates along the and ML axis of wt and *cdh1* embryos (n = 3 embryos) as a function of time during gastrulation (plotted from 7.75–9.5 hpf in 15 min intervals). Negative values represent compression. Mean ± s.e.m.

## DISCUSSION

Our findings highlight a new multi-tiered mechanical mechanism essential for regulating global morphogenetic flows in the ANP by a coordinated spatiotemporal-dependent interplay of extrinsic forces and mechanical tissue coupling. Inter-tissue adhesion and force coupling are thus emerging as key processes in shaping developing embryos ^16, 31, 48–50^. We propose that tissue flows in the ANP are critical to regulate the final tissue shape at the end of gastrulation preceding neurulation. Tissue flows were previously shown to depend on an interplay of multiple tissue intrinsic and/or extrinsic processes which can provide forces necessary to drive oriented tissue flows in the embryo ^51–56^. Interestingly, we found that forces exerted on neuroectoderm cells primarily impact on their movements and less on cell shape changes, suggesting a high degree of tissue fluidity, which has recently been identified as a key mechanical property in reshaping tissues ^13, 57–59^.

Our combined experimental and theoretical results suggest mechanical transitions at different developmental times. Prior to internalisation, forces directed towards the posterior are likely to be generated by pulling (traction) forces of migrating mesendoderm front cells on the overlying neuroectoderm. Our model implies that only the front of the mesendoderm is actively migrating which agrees with our experimental observations. In contrast, long-lasting inter-tissue adhesion of mesendoderm rear cells enables opposing anterior-directed dragging of neuroectoderm cells. Asymmetric distribution of E-cadherin in the axial mesendoderm seems to play a key role in this process. Interestingly, Protocadherin-18a was shown to be enriched as well in the posterior axial mesendoderm where it regulates E-cadherin levels ^60^. These observations raise the intriguing possibility that the mesendoderm collective is composed of different cell populations with distinct force-generating abilities, which is in agreement with recent reports in zebrafish showing that front and rear cells of the migrating mesendoderm, and also of the lateral line primordium, have specialised functions ^61, 62^. We propose that observed differential behaviour of mesendoderm cells can lead to local mechanical instabilities in the ANP that initiate tissue folding and internalisation. During tissue folding, out of plane forces (represented by a sink in the model) drive convergence in the ANP, which we infer as stresses generated by continuous internalisation of cells. This is accompanied by two anterior-directed forces, which are likely to be produced by mesendoderm migration and pushing forces due to AP axis extension of posterior tissues, such as the notochord undergoing CE. This accords with observations in the *Xenopus* neural tube where cell intercalation driven CE occurs primarily in the posterior region and pushes the anterior neural plate forward ^14^. Notably, our findings indicate that convergence in the anterior neural plate is mechanistically different to previously described types that are mainly driven by polarised cell intercalations ^7^.

Recent studies showed that diverse specialised cellular mechanisms control tissue folding in various organisms ^63–68^. Our findings reveal new mechanistic insights into tissue folding by favouring a multi-tiered inter-tissue adhesion model regulating the earliest event of folding in the prospective forebrain. This differs from previously described mechanisms driving neural plate folding during neural tube formation in neurulation. Neural tube morphogenesis in vertebrates is known to be regulated by actomyosin driven apical cell constriction, polarised cell intercalations of converging cells leading to axis extension and contribution of apoptotic cells, which together generate intrinsic forces that drive formation of hinge points and bending of the neural plate ^21, 69–73^. However, recent work from mammalian spinal neural tube folding indicated that hinge points emerge passively in response to external forces ^74^.

Future work will be needed to understand how tissue morphogenesis feeds back into tissue patterning of the emerging forebrain, and in particular how morphogen signalling, regional cell fate specification and robust boundary formation can be established in a dynamically rearranging tissue.

## Acknowledgements

We are thankful to all members of our labs for continuous support and comments on the manuscript. We are grateful to the Warwick BSU aquatics facility for zebrafish care and CAMDU (Computing and Advanced Microscopy Unit) for their support and assistance in this work. This work was supported by the Medical Research Council (MRC) Doctoral Training Partnership (MR/N014294) to A.I., the Biotechnology and Biological Sciences Research Council (BBSRC) MIB Doctoral Training Partnership (BB/T00746X/1) to E.S., an EPSRC grant award (EP/V062522/1) to T.B., the Max Planck Society to P.A.H. and a BBSRC grant award (BB/T016492/1) and the Warwick Quantitative Biomedicine Programme funded by the Wellcome Trust Institutional Strategic Support Fund (ISSF) to M.S.

## Author Contributions

A.I. and M.S. conceived the project and designed the research. A.I. and E.S performed the experiments and analysed the data. J.E.L. and T.B. performed computational image processing and analysis. P.A.H. performed the numerical simulations and modelling. M.T. provided reagents and conceptual input. A.I. and M.S. wrote the manuscript. All authors edited the manuscript.

---

**Supplementary Fig. 1.**
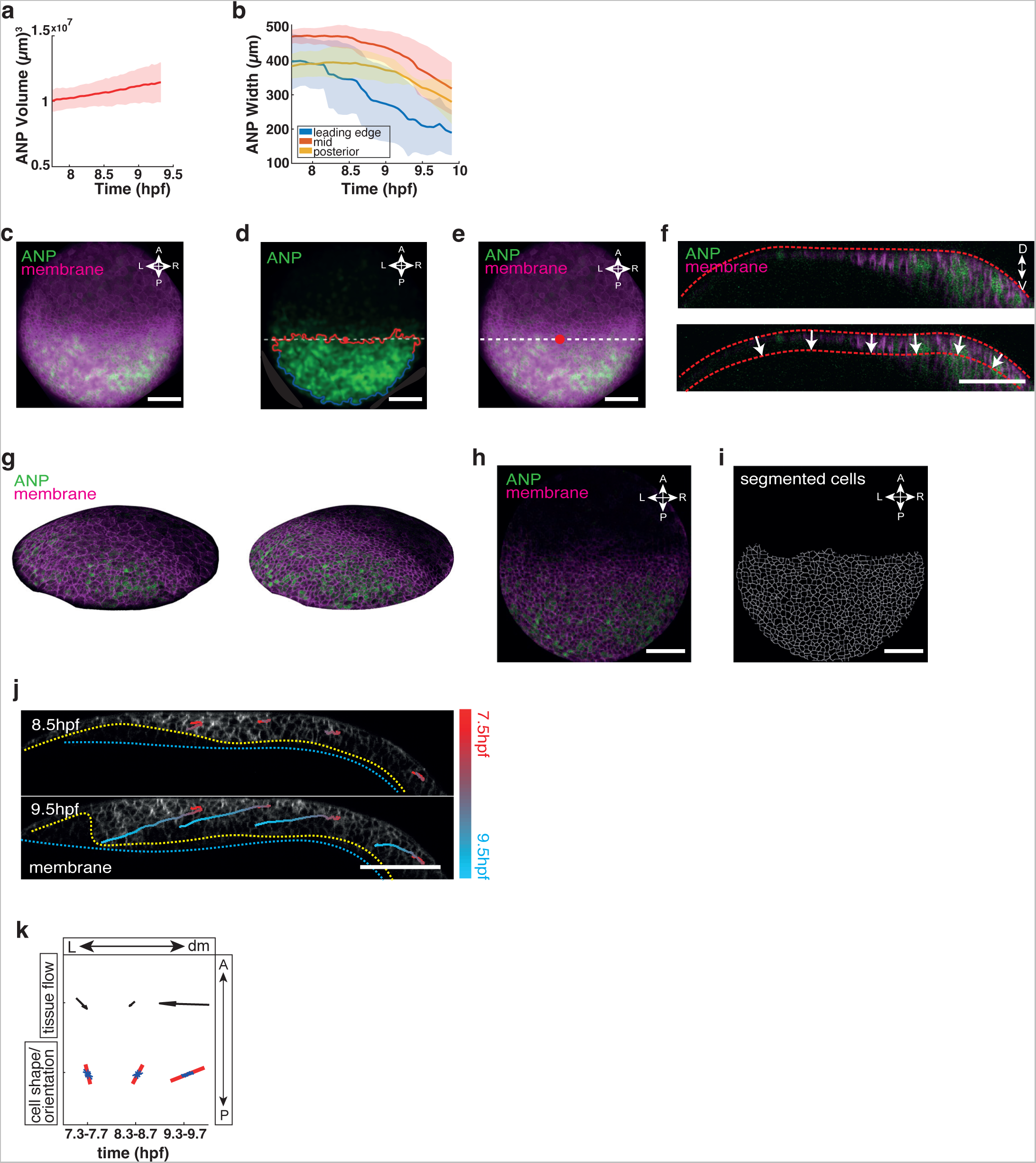
Image processing and analysis of anterior neural plate tissue movements and shape changes during gastrulation. **a)** Anterior neural plate (ANP) tissue volume in wild type (wt) embryos over time of gastrulation (hours post fertilisation, hpf) (n = 3 embryos). Mean ± s.e.m. **b)** ANP width at different locations within the tissue from anterior to posterior edge in wt embryos over time of gastrulation (n = 3 embryos). Mean ± s.e.m. **c)** Mean intensity projection of membrane marker (mRFP) and ANP tissue marker (otx2:Venus) in wt embryos. Animal pole view; anterior (A), posterior (P), left (L) and right (R) axes indicated. Scale bar, 100µm. **d)** Identifying the anterior leading edge of the otx2:Venus region as the approximately linear portion (red) of the contour surrounding the region, with the origin (red circle) placed at the centre of mass of the region projected onto the linear approximation (white dashed line) of the leading edge. View and orientation as in (c). Scale bar, 100µm. **e)** The projection in (c) indicating the ANP leading edge (white dashed line) and the intersection of the AP tissue axis with the leading edge (red circle). Embryo labelled with membrane marker (mRFP) and ANP tissue marker (otx2:Venus). View and orientation as in (c). Scale bar, 100µm. **f)** Segmented surface (red line) of the embryo is translated inwards along the surface normal (white arrows), allowing the mapping of points on the interior into the translated surface (inner red line). Sagittal sections, anterior to the left. Dorsal (D) up and ventral (V) down. Scale bar, 100µm. **g)** Surfaces with fluorescence markers [as in (c)] mapped from the image at two locations, moving inwards from left to right. **h)** Surface fluorescence values from (g) mapped onto the plane. Fluorescent markers, view and orientation as in (c). Scale bar, 100µm. **i)** Segmented cells from surface projection shown in (h). Fluorescent markers, view and orientation as in (c). Scale bar, 100µm. **j)** Confocal images (sagittal sections) showing representative cells tracked within the ANP in wt embryos through gastrulation. Dorsal up, ventral down, anterior to the left and posterior to the right. Cell membranes (mRFP, white) labelled. Yellow line outlines neuroectoderm to mesendoderm border, blue line indicates mesendoderm/yolk interface. Colour code: cells tracked from 7.5hpf (red) to 9.5hpf (blue). Scale bar, 100µm. **k)** Orientation of tissue flow versus cell orientation and shape within cell domains from the right half of the ANP tissue. Arrows indicate the average direction of tissue flows over 20 minutes either side of the indicated time point. Red lines indicate average cell shape/orientation within this time frame. Blue lines represent a histogram of orientations of individual cells (n= 3 embryos).

**Supplementary Fig 2.**
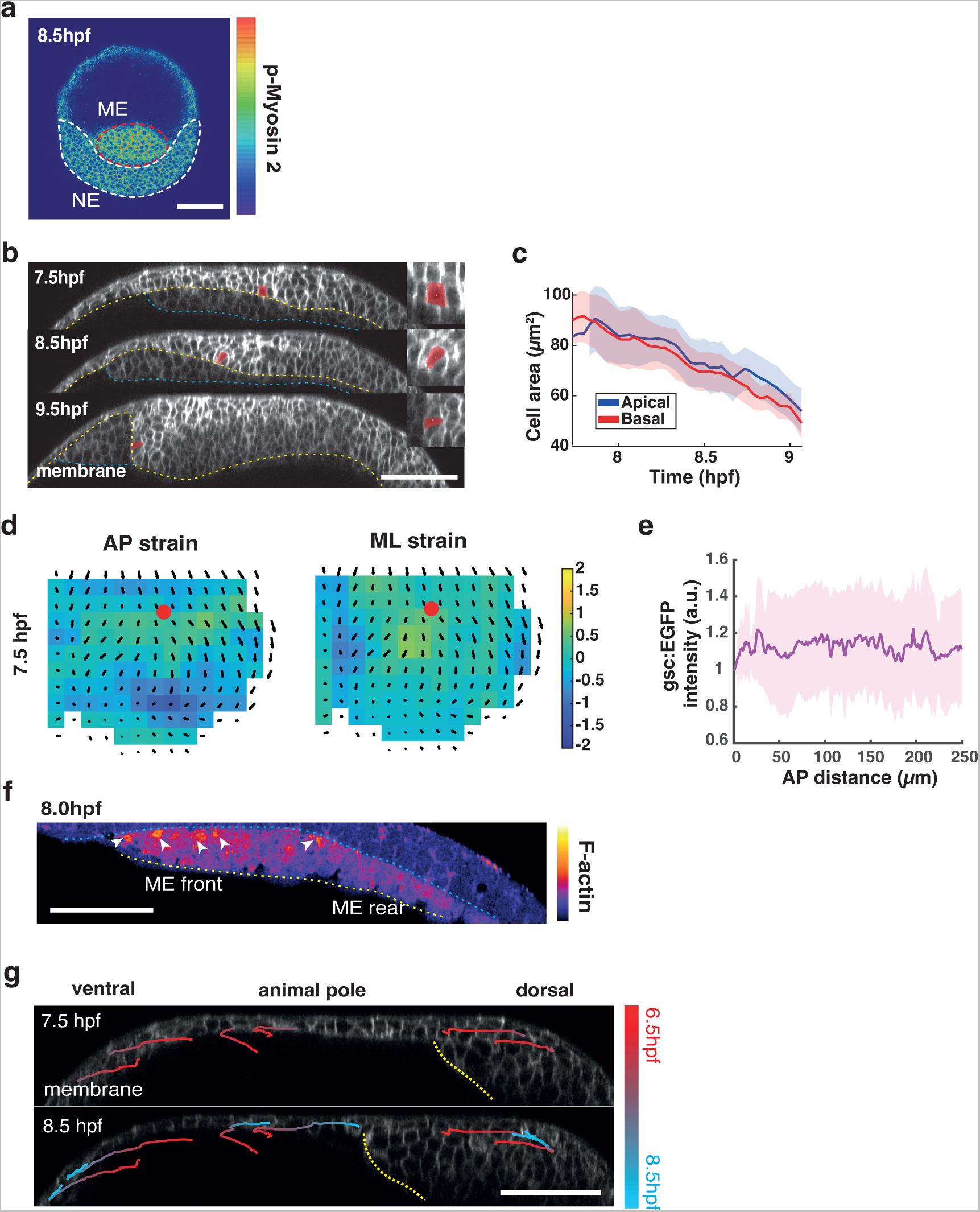
Heterotypic tissue interactions drive ANP tissue folding and neuroectoderm internalisation. **a)** Whole mount staining of phospho-myosin 2 in wild type (wt) embryos. White area demarcates neuroectoderm (NE) cells in the anterior neural plate (ANP), red area indicates mesendoderm (ME). Scale bar = 100µm. **b)** Confocal images (sagittal sections) representing the anterior neural plate (ANP) in wild type (wt) embryos through gastrulation (7.5/8.5/9.5 hours post fertilisation, hpf). Dorsal up, ventral down, anterior left and posterior right. Cell membranes (mRFP, white) labelled. Yellow line outlines neuroectoderm ventral and anterior border to mesendoderm, blue line indicates mesendoderm/yolk interface and white line marks non-neuroectodermal tissue. Representative internalising cell highlighted in red illustrates shape changes over time. Scale bar, 100µm. **c)** Apical (blue) and basal (red) cell surface area of cells around the time of internalisation (n = 3 embryos). Mean ± s.e.m. **d)** Time-averaged ANP domain strains along the anterior-posterior (AP) and mediolateral (ML) axes of wild type embryos (n = 3 embryos). Average normal strain rate is colour-coded (minimum, green (0); maximum stretch, yellow; maximum compression, dark blue). Time-averaged tissue flows are indicated. Red dot marks intersection of ANP leading edge with AP axis. **e)** Intensity levels of gsc:EGFP (a.u.) along the front-rear (AP) axis (0µm = leading edge) of mesendoderm cells. **f)** Confocal images (sagittal sections) of F-actin (lifeact:GFP) in the mesendoderm tissue (8 hpf). F-actin enrichment at protrusions at the neuroectoderm interface indicated (white arrowheads). Scale bar 100 µm. **g)** Confocal images (sagittal sections) with time-resolved neuroectoderm cell tracks within the ANP around the animal pole in wt embryos. Cell membranes (mRFP, white) labelled. Orientation as in (b). Leading edge of the mesendoderm approaching the animal pole (AP) indicated (yellow dotted line). Colour code: 6.5hpf (red) to 8.5hpf (blue). Scale bar, 100µm.

**Supplementary Fig. 3.**
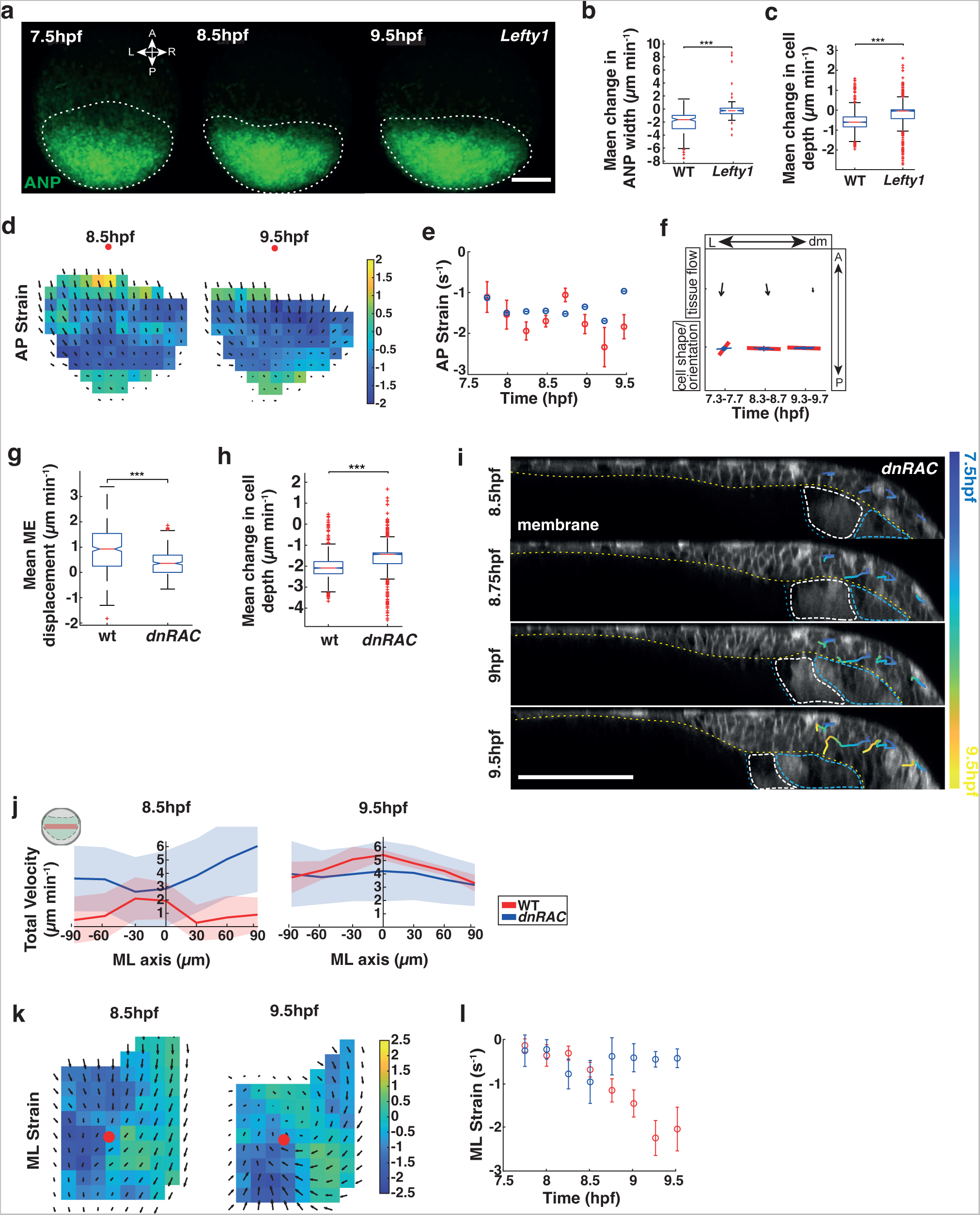
Mesendoderm is necessary for anterior neural plate folding and internalisation. **a)** Live imaging of neuroectoderm cells (maximum projections) in the anterior neural plate (ANP) of *Tg(otx2:Venus) lefty1* morphant embryos during gastrulation (7.5-9.5 hours post fertilisation, hpf). White dotted line outlines the ANP. Scale bar, 100µm. **b)** Mean width change per minute of ANP tissue in *lefty1* morphant and wt embryos (n = 3 embryos). Kruskal-Wallis test (***, p<0.005). Mean ± s.e.m. **c)** Mean neuroectoderm depth change per minute in *lefty1* morphant and wt embryos (n = 3 embryos). Kruskal-Wallis test (***, p<0.005). Mean ± s.e.m. **d)** Time-averaged ANP domain strains along the anterior-posterior (AP) axis of *lefty1* embryos (n = 3 embryos). Average normal strain rate is colour-coded (minimum, green (0); maximum stretch, yellow; maximum compression, dark blue). Time-averaged tissue flows are indicated. Red dot marks intersection of ANP leading edge with AP axis. **e)** Maximum absolute strain rates along the and AP axis of wt and *lefty1* embryos (n = 3 embryos) as a function of time during gastrulation (plotted from 7.75–9.5hpf in 15 min intervals); amount of stretch/compression within each sector is plotted along the y axis. Mean ± s.e.m. **f)** Orientation of tissue flow versus cell orientation and shape within cell domains in *lefty1* morphant embryos (n= 3 embryos). Arrows indicate the average direction of tissue flows over 20 minutes either side of the indicated time point. Red lines indicate average cell shape/orientation within this time frame. Blue lines represent a histogram of orientations of individual cells. **g)** Mean displacement change per minute of the mesendoderm (ME) leading edge in *dnRAC1* and wt embryos (n = 3 embryos). Kruskal-Wallis test (***, p<0.005). Mean ± s.e.m. **h)** Mean neuroectoderm depth change per minute of ANP cells in *dnRAC1* and wt embryos (n = 3 embryos). Kruskal-Wallis test (***, p<0.005). Mean ± s.e.m. **i)** Confocal images (sagittal sections) with time-resolved neuroectoderm cell tracks in the ANP of host embryos transplanted with *dnRac1* ME cells through gastrulation. Dorsal up, ventral down, anterior left and posterior right. Cell membranes (mRFP, white) labelled. Yellow dotted line outlines neuroectoderm/mesendoderm border, blue line indicates host ME, transplanted ME cells outlined with white. Colour code: 7.5hpf (blue) to 9.5hpf (yellow). Scale bar, 100µm. **j)** Total velocities at 8.5hpf and 9.5hpf along the mediolateral (ML) axis of *dnRAC1* embryos (n = 3 embryos). X-axis: 0 marks the dorsal midline; negative, left and positive, right. Mean ± s.e.m. **k)** Time-averaged ANP domain strains along the medio-lateral (ML) axis of *dnRAC1* embryos (n = 3 embryos). Average normal strain rate is colour-coded (minimum, green (0); maximum stretch, yellow; maximum compression, dark blue). Time-averaged tissue flows are indicated. Red dot marks intersection of ANP leading edge with AP axis. **l)** Maximum absolute strain rates along the ANP midline along the ML axis of wt and *dnRAC1* (n = 3 embryos) embryos as a function of time during gastrulation (plotted from 7.75–9.5hpf in 15 min intervals). Amount of stretch/compression within each sector is plotted along the y axis. Mean ± s.e.m.

**Supplementary Fig. 4.**
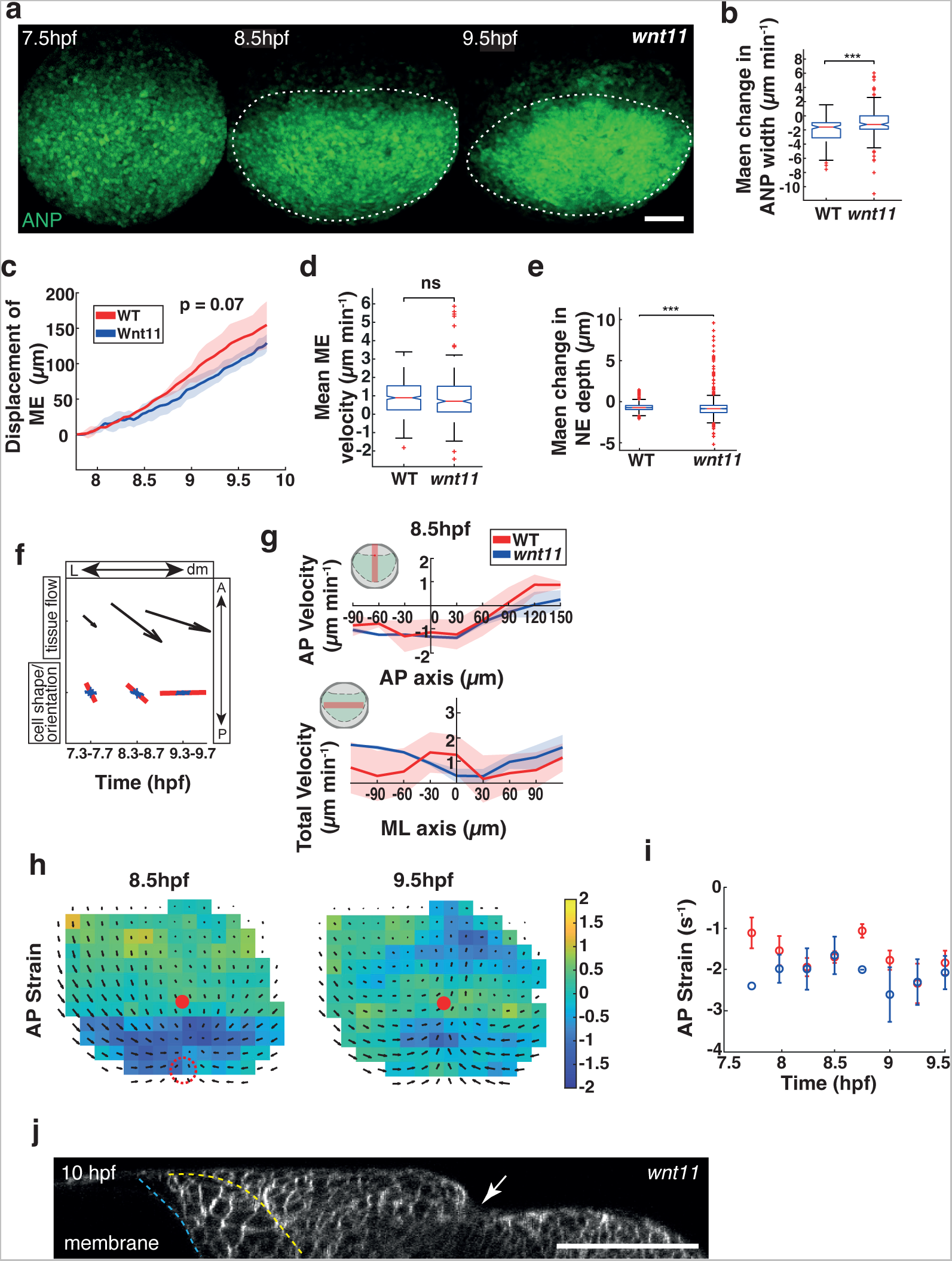
Convergent extension is required for anterior neural plate tissue shape by driving ventral tissue extension. **a)** Live imaging of neuroectoderm cells (maximum projections) in the ANP of *Tg(otx2:Venus); wnt11* mutant embryos during gastrulation (7.5-9.5 hours post fertilisation, hpf). White dotted line outlines the ANP. Scale bar, 100µm. **b)** Mean width change per minute of ANP tissue in *wnt11* mutant and wt embryos (n = 3 embryos). Kruskal-Wallis test (***, p<0.005). Mean ± s.e.m. **c)** Anterior displacement of mesendoderm (ME) cells in *wnt11* mutant and wt embryos (n = 3 embryos). Mean ± s.e.m. **d)** Mean displacement change per minute of leading edge ME cells in *wnt11* mutant and wt embryos (n = 3 embryos). Kruskal-Wallis test (ns, p>0.05). **e)** Mean depth change of neuroectoderm (NE) cells per minute in *wnt11* mutant and wt embryos (n = 3 embryos). Kruskal-Wallis test (***, p<0.005). **f)** Orientation of tissue flow versus cell orientation and shape within cell domains in *wnt11* mutant embryos (n= 3 embryos). Arrows indicate the average direction of tissue flows over 20 minutes either side of the indicated time point. Red lines indicate average cell shape/orientation within this time frame. Blue lines represent a histogram of orientations of individual cells. Anterior (A), posterior (P), left side (L) and dorsal midline (dm) indicated. **g)** Cell velocities at 8.5hpf in anterior-posterior (AP) direction along the dorsal midline and total velocities along the mediolateral (ML) axis of the embryo in wt and *wnt11* embryos (n = 3 embryos). AP velocity, X-axis: 0 marks anterior edge of the ANP; negative, positioned anterior and positive, positioned posterior. Y-axis: negative, posterior and positive, anterior-directed flows. Total velocity, X-axis: 0 marks the dorsal midline; negative, left and positive, right. Mean ± s.e.m. **h)** Time-averaged ANP domain strains along the anterior-posterior (AP) axis of *wnt11* embryos (n = 3 embryos). Average normal strain rate is colour-coded (minimum, green (0); maximum stretch, yellow; maximum compression, dark blue). Time-averaged tissue flows are indicated. Red dot marks intersection of ANP leading edge with AP axis. Dotted circle depicts internalisation site. **i)** Maximum absolute strain rates along the and AP axis of wt and *wnt11* embryos (n = 3 embryos) as a function of time during gastrulation (plotted from 7.75–9.5hpf in 15 min intervals). Negative values indicate compression. Mean ± s.e.m. **j)** Confocal image (sagittal section) indicating a dorsal kink in the ANP of *wnt11* morphant embryos at 10 hpf. Dorsal up, ventral down, anterior to the left and posterior to the right. Cell membranes (mRFP, white) labelled. Yellow line marks interface between neuroectoderm and mesendoderm, blue line outlines mesendoderm-yolk interface. Scale bar 100µm.

**Supplementary Fig. 5.**
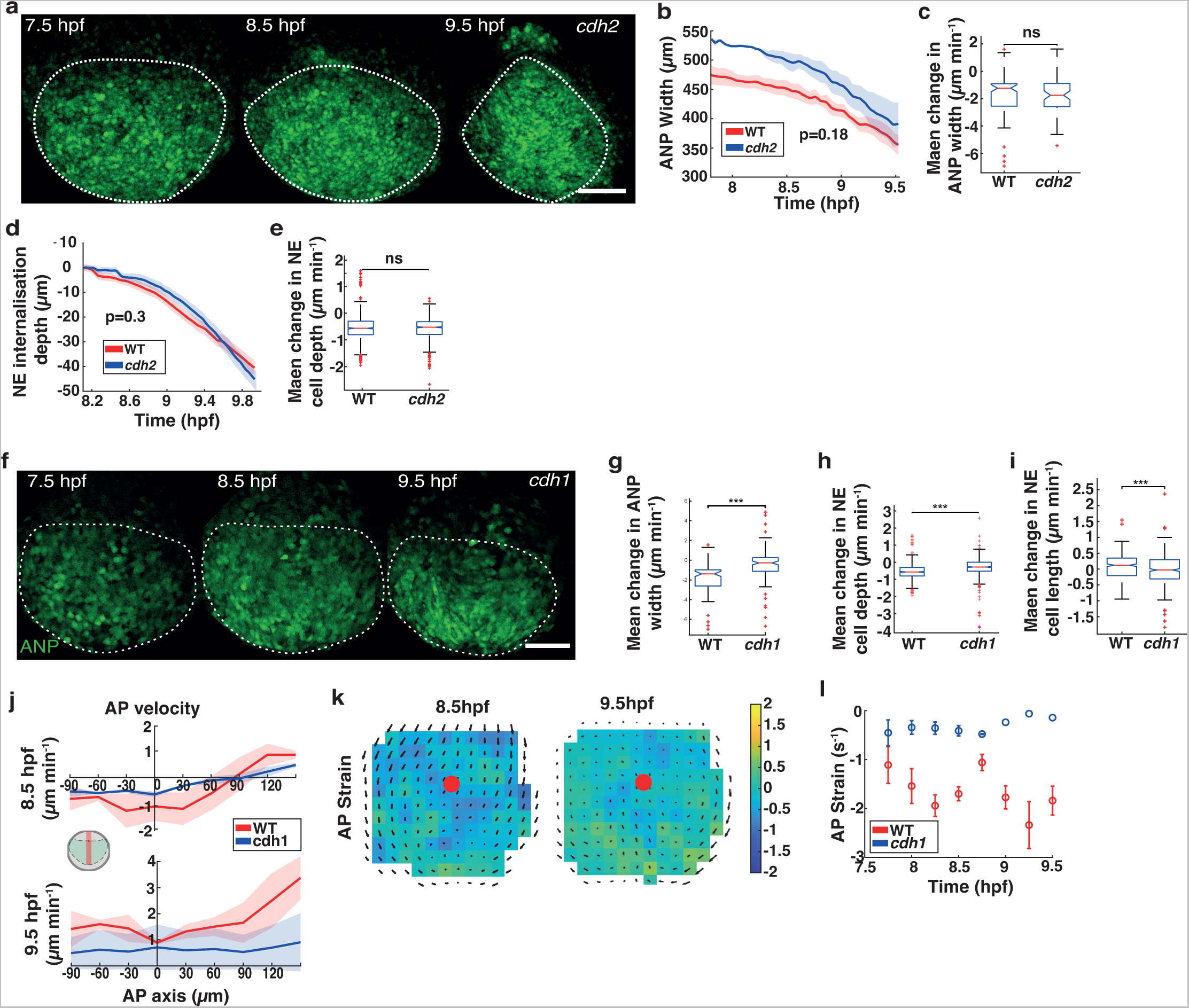
E-cadherin-mediated inter-tissue interaction is required for neuroectoderm internalisation. **a)** Live imaging of neuroectoderm cells (maximum projections) in the anterior neural plate (ANP) of *Tg(otx2:Venus) cadherin 2 (cdh2)* morphant embryos during gastrulation (7.5-9.5 hours post fertilisation, hpf). White dotted line outlines the ANP. Scale bar, 100µm. **b)** ANP tissue width in wildtype (wt, red) and *cdh2* morphant (blue) embryos over time of gastrulation (hours post fertilisation, hpf) (n = 3 embryos). Mean ± s.e.m. **c)** Mean width change per minute of ANP tissue in *cdh2* mutant and wt embryos (n = 3 embryos). Kruskal-Wallis test (ns, p<0.005). Mean ± s.e.m. **d)** Neuroectoderm (NE) internalisation depth in in wildtype (wt, red) and *cdh2* morphant (blue) embryos over time of gastrulation (hpf) (n=3 embryos). **e)** Mean neuroectoderm depth change per minute over late gastrulation in wt (red) and *cdh2* morphant (blue) embryos (n = 3 embryos). Kruskal-Wallis test (ns, p<0.005). Mean ± s.e.m. **f)** Live imaging of neuroectoderm cells (maximum projections) in the ANP of *Tg(otx2:Venus) cadherin-1 (cdh1)* morphant embryos during gastrulation (7.5-9.5 hours post fertilisation, hpf). White dotted line outlines the ANP. Scale bar, 100µm. **g)** Mean width change per minute of ANP tissue in *cdh1* mutant and wt embryos (n = 3 embryos). Kruskal-Wallis test (***, p<0.005). Mean ± s.e.m. **h)** Mean neuroectoderm (NE) depth change per minute of ANP cells in *cdh1* mutant and wt embryos (n = 3 embryos). Kruskal-Wallis test (***, p<0.005). **i)** Mean length change in NE cell length during internalisation in *cdh1* mutant and wt embryos (n = 3 embryos). Kruskal-Wallis test (***, p<0.005). Mean ± s.e.m. **j)** Cell velocities along the anterior-posterior (AP) axis of wt (red) and *cdh1* (blue) embryos at 8.5 and 9.5 hpf (n = 3 embryos). Y-axis: cell velocity (µm/min); X-axis (µm): 0 marks the dorsal midline of the embryo. Mean ± s.e.m. **k)** Time-averaged ANP domain strain rates along the anterior-posterior (AP) axis of *cdh1* embryos (n = 3 embryos). Average normal strain rate is colour-coded (minimum, green (0); maximum stretch, yellow; maximum compression, dark blue). Time-averaged tissue flows are indicated. Red dot marks intersection of ANP leading edge with AP axis. **l)** Maximum absolute strain rates along the ANP midline along the AP axis of wt and *cdh1* embryos (n = 3 embryos) as a function of time during gastrulation (plotted from 7.75–9.5hpf in 15 min intervals; n=3 embryos). Negative values indicate compression. Mean ± s.e.m.

## Supplementary Videos

**Supplementary Video 1. Live imaging of anterior neural plate dynamics in wild type embryo.**

Time-lapse imaging of wild type (wt) *Tg(otx2:Venus)* embryo injected with membrane RFP (mRFP) marker. Neuroectoderm cells (green) and cell outlines (magenta) are imaged from 7.5 to 10 hpf (192 minutes). Animal pole view; anterior up and posterior down.

**Supplementary Video 2. Live imaging of anterior neural plate folding and neuroectoderm internalisation in wild type embryo.**

Time-lapse imaging of wild type (wt) *Tg(otx2:Venus)* embryo injected with membrane RFP (mRFP) marker. Neuroectoderm cells (green) and cell outlines (magenta) are imaged from 7.5 to 10 hpf (192 minutes). Sagittal section; anterior left, posterior right; dorsal up and ventral right.

**Supplementary Video 3. Live imaging of anterior neural plate dynamics in *lefty 1* morphant embryo.**

Time-lapse imaging of *Tg(otx2:Venus)* embryo injected with *lefty 1* morpholino and membrane RFP (mRFP) marker. Neuroectoderm cells (green) and cell outlines (magenta) are imaged from 7.5 to 10 hpf (185 minutes). Animal pole view; anterior up and posterior down.

**Supplementary Video 4. Live imaging of neuroectoderm internalisation in *wnt11* mutant embryo.**

Time-lapse imaging of *Tg(otx2:Venus)*; *wnt 11* mutant embryo injected with membrane RFP (mRFP) marker. Neuroectoderm cells (green) and cell outlines (magenta) are imaged from 7.5 to 9.5 hpf (144 minutes). Sagittal section; anterior left, posterior right; dorsal up and ventral right.

**Supplementary Video 5. Live imaging of neuroectoderm internalisation in *cadherin-1* morphant embryo.**

Time-lapse imaging of *Tg(otx2:Venus)* embryo injected with *cadherin-1* morpholino and membrane RFP (mRFP) marker. Neuroectoderm cells (green) and cell outlines (magenta) are imaged from 7.5 to 9.5 hpf (124 minutes). Sagittal section; anterior left, posterior right; dorsal up and ventral right.

## Methods

### Fish lines and maintenance

Wild type (wt) AB and TU strains; transgenic lines: *Tg(otx2:Venus)*^32^, *Tg(actb1:lifeact-GFP)* ^74^, *Tg(1.8gsc:GFP-CAAX)* ^31^, *Tg(1.8gsc:GFP)* ^75^. Transgenic mutant homozygous line *Tg(otx2:Venus); wnt11* was generated in-house by crossing *Tg(otx2:Venus)* with *wnt11(silberblick)* mutants ^44^. To generate the *Tg(gsc:Cdh1-EGFP)* line, mouse E-cadherin-GFP-GI-polyA ^46^ was placed downstream of the goosecoid (gsc) promoter in the Tol2-basic vector ^31, 76^.

Zebrafish were maintained as described previously^77^ and embryos were raised at 28-31 °C in E3 buffer. Embryos were selected for imaging using established morphological criteria ^78^. Adult zebrafish were maintained in the University of Warwick aquatics facility, in compliance with the University of Warwick animal welfare and ethical review board (AWERB) and the UK home office animal welfare regulations.

### Microinjections of capped mRNA and morpholino antisense oligonucleotides

Capped mRNA used for injection was synthesised using SP6 mMessage mMachineKit (Ambion). For ubiquitous mRNA expression, 100 pg of *mRFP* mRNA 100 pg of *Lefty1* mRNA ^79^, 75 pg of *h2afva-tagBFP* mRNA ^80^ and 400 pg of *dnRAC1* mRNA ^41^ were injected into one cell stage embryos. Morpholino oligonucleotides (MOs) were designed and synthesized by GeneTools and injected into one-cell stage embryos. To down regulate E-cadherin or N-cadherin, 3-4 ng *cadherin 1* ^31^ or 3 ng *cadherin 2* ^81^ MO was injected into one-cell-stage embryos, respectively. To generate mesoderm (prechordal plate) progenitors, 100 pg *squint* mRNA and 2 ng *sox32/casanova* MO^82^ were injected into one cell stage embryos.

### Sample preparation for live cell imaging

Dechorionated embryos were mounted in 0.8% low-melting-point (LMP) agarose (Invitrogen) into agarose moulds inside a glass fluorodish (WPI World Precision Instrument) and covered with E3 medium with the animal pole of the embryo facing towards the objective. Wild type embryos were checked for normal development and mutant embryos for respective phenotype after imaging (12-24 hours post fertilisation).

### Confocal Imaging

I*n vivo* live imaging and fixed whole mount imaging of embryos were performed using a Zeiss 880 confocal microscope using a Zeiss 25x oil objective (NA = 0.8). The temperature during imaging was kept at 29.5 °C using a temperature chamber. Embryos were mounted at 7.5 hours post fertilisation (hpf) and imaged through to the end of gastrulation (10 hpf). Venus and GFP fluorophores were imaged at 488 nm and RFP at 564 nm, respectively. Z-stacks were taken at a spacing of 1.2 μm with a XY resolution of 0.78 μm per pixel. Images were taken every 180–250 seconds.

### Multiphoton imaging

For *in vivo* live imaging, embryos were imaged using an Olympus FVMPE-RS inverted multiphoton microscope using a 30x silicon objective (NA = 1.05). Proteins were imaged at the following wavelengths: mRFP - 1050 nm; Venus and GFP - 925 nm; BFP - 780 nm, using a Spectra-Physics InSight X3 and MaiTai HP DS-OL laser. Power was adjusted dependent on tissue depth and Z-spacing was set to 1.2 μm and a resolution of 0.53 μm per pixel. Images were taken every 180–220 seconds. Embryos were imaged at 29.5 °C.

### Whole-mount immunohistochemistry and antibodies

For whole-mount immunohistochemistry, embryos were fixed overnight at 70% (7.5 hpf), 80% (8.25 hpf), 90% (9 hpf) and 100% (10 hpf) in 4% paraformaldehyde in 1x PBS. After fixation they were washed 6 times in PBS with 0.1% Triton-X in 1x TBS (TBST) then permeabilised with 0.5% Triton-X in 1xTBS. Embryos were subsequently blocked in 0.1% Triton-X and 5% goat serum in 1xTBS. Phosphorylated myosin II was detected using a primary rabbit anti-phosphomyosin antibody (Sigma-Aldrich, F3648; 1/200 dilution). Incubation with primary antibodies was performed overnight at 4°C in TBS-T containing 0.1% Triton-X and 5% goat serum in 1xTBS at 4°C. Embryos were consequently washed with TBS-T six times for 10 min each and incubated overnight with secondary antibody (Alexa 488-conjugated goat anti-rabbit, ThermoFisher Scientific, A-11008; 1/5000 dilution) and rhodamine-phalloidin for F-actin staining (ThermoFisher Scientific, R415; 1/200 dilution). Embryos were washed three times for 5 min with TBST and nuclei were stained with DAPI nuclei acid stain (ThermoFisher Scientific, D1306) for 30 minutes before a final round of three times 5 minute washes in TBS-T.

### Transplantation assays

For cell transplantation experiments, donor and host embryos were kept in Danieau’s solution (58 mM NaCl, 0.7 mM KCl, 0.4 mM MgSO_4_, 0.6 mM Ca(NO_3_)_2_, 5mM HEPES pH 7.6) after dechorionation. To produce mesendoderm (prechordal plate) progenitors, *Tg(gsc:GFP-CAAX)* donor embryos injected at one-cell stage with 100 pg of *squint* mRNA, 400pg of *dnRAC1* mRNA and 2 ng morpholino against *sox32/casanova*. Host *Tg(1.8gsc:EGFP-CAAX)* embryos were injected at one-cell stage with 50 pg of *mRFP* mRNA. Groups of mesoderm-induced cells (100–200 cells) were then removed from the animal pole of donor embryos at sphere stage using a glass transplantation needle (20 µm diameter) and transplanted below the neuroectoderm cells in front of the endogenous prechordal plate of host embryos between 7 hpf and 8 hpf (before ANP internalisation). Transplanted embryos were mounted for imaging as described above.

### Image processing

#### Rotation and origin

The otx2:Venus channel was used to identify the angle of rotation about the *z*-axis and identify the (*x*,*y*)-coordinates of the new origin, with the aim of aligning the front of the otx2:Venus region with the *x*-axis, placing the new origin at the centre of the anterior edge (Supplementary Fig. 1c-e). The time point corresponding to tissue internalisation was selected to compute these values as it represents the time when the anterior edge of the tissue is flat. The mean intensity *z*-projection of the channel was taken and Otsu thresholding applied. The contour around the largest connected component of this thresholded image was taken to be the boundary of the otx2:Venus region. Lines between all points of the contour with length between 20 and 200 pixels were drawn to identify the modal unsigned direction, represented as the angle from the positive *x*-axis between 0 and π radians, using a bin width of π/10. All lines with an angle not in the modal or adjacent histogram bins were discarded, and the remaining lines were used to group points on the contour by connectivity. The largest set of nodes connected by these lines was taken as the points along the front (Supplementary Fig. 1d, red line). Points along the front were sorted by position along the contour, starting at the end of the front that preserves the handedness of the axes in the following step. The average signed direction angle of all remaining lines connecting points on the front to later points in the sorted sequence was taken to be the new positive *x*-direction, *v*, and the new positive *y*-direction, *u*, was taken as perpendicular to this direction, pointing away from the otx2:Venus region the front. The origin in these new coordinates was given by the centre of mass of the otx2:Venus region in the direction *v*, and the centre of the points on the front in the direction *u*.

#### Surface projection

The whole embryo was segmented by applying Gaussian smoothing (*s* = 3 pixels) to the membrane channel followed by adaptive Phansalkar thresholding (radius = 30 voxels) ^83^ and selecting the largest connected component of the binary output as the segmented embryo. Points on the outer surface (Supplementary Fig. 1f) of the embryo were identified from the voxels with maximal *z*-component for each (*x*,*y*)-coordinate covered by the segmented embryo. For images with the outer surface facing *z* = 0, the minimal *z*-component was used. This orientation was detected by first computing the mean distance to the origin, defined above, of all pixels in the segmented embryo in each *z*-slice and taking the median of these values in the top and bottom halves of the *z*-stack. The half with the smaller median was taken to contain the outer surface, since it represents the narrower portion of the embryo in the imaged volume. The *z*-coordinate of the new origin was taken to be the *z*-coordinate of the surface point nearest the origin in the (*x*,*y*)-plane identified in the previous section at the same time point. All surface points were translated to place the new origin at (0,0,0). Each translated surface point, *p* = (*x*,*y*,*z*), was mapped to the point *q*=*r*(*p*)(*x*,*y*), with scale factor *r*(*p*) such that *q* and *p* have the same length. An inverse mapping from integer (*x*,*y*)-coordinates onto the surface was found by linear interpolation of *z* from projected surface points followed by Gaussian smoothing of *z*-values (*s* = 20 pixels), with the mapping completed by rescaling the (*x*,*y*)-coordinates of the surface points to maintain lengths as in the forward mapping. This yielded a smooth surface and a corresponding planar mapping to a 2D image. Volumetric mapping was achieved by translating surface points along the surface normal in steps equal to the original *z*-resolution (Supplementary Fig. 1f), yielding a stack of projected 2D images with fluorescence values mapped using nearest neighbour interpolation (Supplementary Fig. 1g-h). This mapping allowed standard image analysis methods to be applied to individual slices without having to contend with multiple layers of the embryo being present within a single slice.

### Image analysis and tissue flows

#### Tissue shape analysis

The shape of the tissue was calculated using the surface projection of the otx2:Venus signal as described above. The projection was then analysed by segmenting the maximum projection using a watershed algorithm in MATLAB. Area of the tissue was calculated using this segmentation. The extent of the left-right axis was calculated by averaging the width of the tissue at five points along the anterior-posterior (AP) axis. All parameters were averaged over different embryos.

#### Analysis of cell shapes

Projected cells were segmented in 2D by first applying Gaussian smoothing (*s* = 3), followed by watershed segmentation of the mRFP channel. The segmented cell shapes were used to measure surface area, orientation and maximum and minimum axis using Matlab’s regionprops function. Values were refactored to account for changes induced by the mapping by taking locations of the extremities of the major and minor axes of the ellipse and projecting these back into ‘original’ coordinates. For calculations of the orientation and dynamics of ‘domains’ of cells, boxes with sides of 30 μm (approx. 15 cells) were used and the average shape of cells within these domains was calculated over a period of 20 minutes. The centre of the domain was centred on an individual cell that was manually tracked from the anterior edge of the ANP. To investigate the dynamics of cells as they internalise, cells were manually backtracked to mid-gastrulation. The apical and basal area of the cell was then measured using ImageJ (Fiji)^84^. The dorso-ventral extent of cells was calculated by the distance between these sections, distances were averaged between multiple cells and multiple embryos.

#### Cell depth during internalisation

To determine the depth of individual neuroectoderm cells in the ANP over the period of gastrulation, cells that were initially within the dorsal cell layer along the dorsal midline of the embryo were manually tracked within multiple embryos. The minimum distance to the surface of the embryo calculated.

#### Quantifying tissue flows

Projected images were analysed using the MATLAB application PIVlab. 30 μm^2^ boxes were created spanning from the origin along the lateral and AP axes. Velocities of cells within these boxes were then averaged over 15 minutes to calculate the average velocity of domains within the tissue.

#### Measuring heterotypic cell-cell interactions

To calculate the contact time between mesendoderm and neuroectoderm cells in the ANP, mesendoderm cells were manually tracked. The time period for each new neuroectodermal cell contact was then averaged over the period of imaging to give a mean time of cell contact. This was completed for a point at the front (anterior), the middle and the rear (posterior, adjacent to the notochord) of the mesendoderm. Contact times were then averaged over different embryos.

#### Quantification of E-cadherin and F-actin levels

Measurements were performed in ImageJ/Fiji using sagittal sections from live imaging of gsc:cdh1-EGFP, gsc:EGFP control or lifeact:EGFP at ~8 hpf (before internalisation) in the embryo through the reslice function. A freehand line was drawn in the mesendoderm close to the neuroectoderm interface, starting from the front to the rear of the collective, covering around 250 μm of tissue. Plot profiles of signal intensity were generated for three different sections per experimental conditions, normalised and averaged over all sections.

### Calculation of tissue strain rates

Strain rates in the neuroectoderm were calculated from the tissue flows using spatial derivatives of velocities in neighbouring domains (~40 µm^2^) ^36^. The strain rates in the anterior-posterior (AP) and left-right (LR) directions are

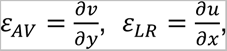

respectively, where *x* and *y* are the coordinates along the LR and AP axes, and *u*, *v* are the velocities in these directions. These strain rates determine the stretch (positive strain rate) or compression (negative strain rate) in these directions.

### Statistical and data analysis

Data was stored in mat files and analysis was performed using MATLAB. All data plots were generated in MATLAB using built-in functions. Box plots generated using the boxplot function; central line indicates the median, and the bottom and top edges of the box indicate the 25th and 75th percentiles, respectively. The whiskers extend to the most extreme data points not considered outliers, and the outliers are plotted individually using the + marker symbol. A one-way Kruskal–Wallis test was used when comparing whether different groups came from the same distribution and implemented using the MATLAB kruskalwallis function.

## Theoretical model

Details of the theoretical model can be found in the Supplemental Note.

## Data and code availability

All data supporting the findings of this study are available from the corresponding authors on reasonable request.

