## supplemental note for "A multi-tiered mechanical mechanism shapes the early neural plate"

This Supplemental Note is divided, like Gaul [S1], into three parts. In the first part, we give the expressions for the flow singularities used to discuss the experimental flow fields qualitatively. In the second part, we derive the expressions for the regularised singularities used to fit the experimental flow fields and give details of the fitting procedure. In the final part, we discuss modelling tissue compressibility and the YSL/EVL margin and extend the discussion of the main text.

### I. FUNDAMENTAL SOLUTIONS OF STOKES FLOW IN TWO DIMENSIONS

In the main text, we analyse the flows in the neurectoderm qualitatively by discussing the flows of a two-dimensional unbounded incompressible viscous fluid resulting from different point forces (representing migration and drag forces exerted by the mesendoderm) and point sinks (representing ingression or the compressibility of the tissue). Since the Stokes equations that describe these flows are linear, they are the superposition of the flows resulting from the individual point forces and point sources [S2]. This Section gives the expressions for these singularities or fundamental solutions.

#### A. The Stokeslet in two dimensions

We consider the incompressible Stokes equations [S2] with a point force $\boldsymbol{f}$ at $\boldsymbol{x} = \boldsymbol{x_0}$. Without loss of generality, we may take $\boldsymbol{x_0} = \boldsymbol{0}$ by translational invariance. The associated flow field is the solution of [S2]

$$\boldsymbol{0} = -\boldsymbol{\nabla} p + \nabla^2 \boldsymbol{u} + \boldsymbol{f}\delta(\boldsymbol{x}), \qquad \boldsymbol{\nabla}\cdot\boldsymbol{u} = 0, \tag{1}$$

in which $\boldsymbol{u}(\boldsymbol{x})$ is velocity, $p(\boldsymbol{x})$ is pressure, and where $\delta$ denotes the Dirac delta function. Here, and throughout this Supplemental Note, we have nondimensionalised the equations in such a way as to set the fluid viscosity equal to $\mu = 1$.

The solution of Eqs. (1) is called the Stokeslet, and, in two spatial dimensions, it is [S2]

$$\boldsymbol{u} = \frac{\boldsymbol{f}}{4\pi}\cdot\left(-\log r\,\boldsymbol{I} + \frac{\boldsymbol{x}\boldsymbol{x}}{r^2}\right), \tag{2}$$

where $r = \|\boldsymbol{x}\|$ and $\boldsymbol{I}$ is the identity. This flow field diverges at $r = 0$ due to the singular nature of the force applied there. Moreover, $\boldsymbol{u}$ diverges unphysically as $r \to \infty$; this is known as Stokes' paradox, and is a limitation of the two-dimensional Stokes equations in an unbounded fluid [S3]. Owing to this divergence, the Stokeslet has stagnation points on the flow axis defined by $\boldsymbol{f}$. These are clearly unphysical [S4], so can be ignored in the qualitative analysis in the main text.

#### B. Sink flow in two dimensions

The flow resulting from a sink of strength $q$ at $\boldsymbol{x} = \boldsymbol{0}$ in incompressible Stokes flow is described by the equations [S2]

$$\boldsymbol{0} = -\boldsymbol{\nabla} p + \nabla^2 \boldsymbol{u}, \qquad \boldsymbol{\nabla}\cdot\boldsymbol{u} = -q\delta(\boldsymbol{x}). \tag{3}$$

The solution of these equations in two dimensions is well-known; it is [S2]

$$\boldsymbol{u} = -\frac{q\boldsymbol{x}}{2\pi r^2}. \tag{4}$$

### II. FITS OF THE EXPERIMENTAL FLOW FIELDS USING REGULARISED SINGULARITIES

For a more quantitative representation of the experimental flows, we must regularise the divergence of the fundamental solutions at the origin, i.e. at the point where the force is applied or the sink is located, and the divergence in the far-field associated with Stokes' paradox.

To regularise Stokes' paradox, we include a friction force representing the friction between the neurectoderm and the EVL. To regularise the divergence at the origin, we smear out the point forces and point sinks. The equations (1) and (3) defining the fundamental solutions are thus replaced with

$$\boldsymbol{0} = -\boldsymbol{\nabla} p + \nabla^2 \boldsymbol{u} - \gamma^2\boldsymbol{u} + \boldsymbol{f}\phi_\varepsilon(\boldsymbol{x}), \qquad \boldsymbol{\nabla}\cdot\boldsymbol{u} = 0 \tag{5}$$

and

$$\boldsymbol{0} = -\boldsymbol{\nabla} p + \nabla^2 \boldsymbol{u} - \gamma^2\boldsymbol{u}, \qquad \boldsymbol{\nabla}\cdot\boldsymbol{u} = -q\phi_\varepsilon(\boldsymbol{x}), \tag{6}$$

respectively. In these equations, $\gamma$ is a friction coefficient, and $\phi_\varepsilon$ is a "blob function", depending on a parameter $\varepsilon$ that represents the characteristic distance over which the point force or point sink is smeared out.

The Stokes equations with a friction term $-\gamma^2\boldsymbol{u}$ added in this way are known as the Brinkman equations and also constitute an extension of Darcy's law for flow in porous media [S5].

Smearing out force singularities using a blob function $\phi_\varepsilon$, as in Eqs. (5) and (6), is an efficient technique [S6–S8] for numerical solution of hydrodynamic equations by regularisation of the singular boundary integral formulation [S9]. In these numerical approaches, the blob function is normalised in such a way that

$$\iint_{\mathbb{R}^2} \phi_\varepsilon(\boldsymbol{x})\,\mathrm{d}^2\boldsymbol{x} = 1, \tag{7}$$

and satisfies $\phi_\varepsilon \longrightarrow \delta$ as $\varepsilon \longrightarrow 0$, but remain finite for all $\varepsilon > 0$, so that the numerical solution for $\varepsilon > 0$ approaches the exact solution as $\varepsilon \longrightarrow 0$. In particular, finite values of $\varepsilon$ do not therefore have a physical meaning, though they can affect, via the choice of blob function, the rate of convergence of the numerical solution to the exact solution [S10].

Here, by contrast, we do ascribe a physical meaning to finite values of $\varepsilon$: they represent the extent over which the mesendoderm exerts friction on the neurectoderm or over which cells in the neurectoderm ingress. We are not aware of a prior use of regularised singularities in this way.

In this context, a two-dimensional "Brinkmanlet" solution of Eqs. (5) was derived in Ref. [S11] for numerical solution of these equations, but the blob function chosen in Ref. [S11] depends on the friction parameter $\gamma$. While this is not an issue for these numerical approaches, it is not appropriate in our case, given the physical meaning of the blob function.

Here, we therefore choose the blob function to be the radially symmetric function

$$\phi_\varepsilon(r) = \frac{r\mathcal{K}_1(r/\varepsilon)}{4\pi\varepsilon^3}, \tag{8a}$$

where we use $\mathcal{K}_0, \mathcal{K}_1, \dots$ to denote the modified Bessel functions of the second kind [S12]. It satisfies the normalisation condition (7). With the hindsight of the calculations that follow below, this somewhat arcane choice of blob function is motivated by the simple form of its Fourier transform [S13]

$$\tilde{\phi}_\varepsilon(k) = \left(1 + \varepsilon^2 k^2\right)^{-2}, \tag{8b}$$

where we have let $k = \|\boldsymbol{k}\|$. This allows the Fourier inversion integrals that arise in the solution of Eqs. (5) and (6) to be computed in closed form.

#### A. Solution of Eqs. (5): the regularised "Brinkmanlet"

We solve Eqs. (5) in Fourier space. Taking their Fourier transform [S13], we find

$$\boldsymbol{0} = -\mathrm{i}\boldsymbol{k}\tilde{p} - k^2\tilde{\boldsymbol{u}} - \gamma^2\tilde{\boldsymbol{u}} + \boldsymbol{f}\tilde{\phi}_\varepsilon(k), \qquad \mathrm{i}\boldsymbol{k}\cdot\tilde{\boldsymbol{u}} = 0, \tag{9}$$

for a radially symmetric blob function, as in Eq. (8a). Dotting the first equation with $\boldsymbol{k}$ and using the second one yields

$$\tilde{p} = \boldsymbol{f}\cdot\left(-\frac{\mathrm{i}\boldsymbol{k}\tilde{\phi}_\varepsilon(k)}{k^2}\right), \tag{10a}$$

and hence, on substituting this into the first of Eqs. (9),

$$\tilde{\boldsymbol{u}} = \boldsymbol{f}\cdot\left[\frac{\tilde{\phi}_\varepsilon(k)}{k^2+\gamma^2}\left(\boldsymbol{I} - \frac{\boldsymbol{kk}}{k^2}\right)\right]. \tag{10b}$$

By symmetry, the regularised "Brinkmanlet" solution is thus

$$\boldsymbol{u} = \boldsymbol{f}\cdot\left[A_\varepsilon(r)\boldsymbol{I} + B_\varepsilon(r)\frac{\boldsymbol{xx}}{r^2}\right], \tag{11}$$

in which the coefficients $A_\varepsilon(r), B_\varepsilon(r)$ remain to be determined by computing the Fourier inversion

$$A_\varepsilon(r)\boldsymbol{I} + B_\varepsilon(r)\frac{\boldsymbol{xx}}{r^2} = \frac{1}{(2\pi)^2}\iint_{\mathbb{R}^2}\frac{\tilde{\phi}_\varepsilon(k)}{k^2+\gamma^2}\left(\boldsymbol{I} - \frac{\boldsymbol{kk}}{k^2}\right)\mathrm{e}^{\mathrm{i}\boldsymbol{k}\cdot\boldsymbol{x}}\,\mathrm{d}^2\boldsymbol{k}. \tag{12}$$

With Cartesian axes in which $\boldsymbol{x} = (r,0)$, $\boldsymbol{k} = k(\cos\theta, \sin\theta)$, taking the trace and the $(1,1)$ component of Eq. (12) respectively, and using results from the Appendix of this Note,

$$2A_\varepsilon(r) + B_\varepsilon(r) = \frac{1}{(2\pi)^2}\int_0^\infty\!\!\int_0^{2\pi}\frac{\tilde{\phi}_\varepsilon(k)}{k^2+\gamma^2}\mathrm{e}^{\mathrm{i}kr\cos\theta}\,k\,\mathrm{d}\phi\,\mathrm{d}k = \frac{1}{2\pi}\int_0^\infty\frac{k\tilde{\phi}_\varepsilon(k)}{k^2+\gamma^2}\mathcal{J}_0(kr)\,\mathrm{d}k, \tag{13a}$$

$$A_\varepsilon(r) + B_\varepsilon(r) = \frac{1}{(2\pi)^2}\int_0^\infty\!\!\int_0^{2\pi}\frac{\tilde{\phi}_\varepsilon(k)}{k^2+\gamma^2}\sin^2\theta\,\mathrm{e}^{\mathrm{i}kr\cos\theta}\,k\,\mathrm{d}\phi\,\mathrm{d}k = \frac{1}{2\pi r}\int_0^\infty\frac{\tilde{\phi}_\varepsilon(k)}{k^2+\gamma^2}\mathcal{J}_1(kr)\,\mathrm{d}k, \tag{13b}$$

where we use $\mathcal{J}_0, \mathcal{J}_1, \dots$ to denote the Bessel functions of the first kind [S12]. We now specialise to the blob function (8a). Noting the partial fraction decomposition [S14]

$$\frac{\tilde{\phi}_\varepsilon(k)}{k^2+\gamma^2} = \frac{1}{\left(\gamma^2\varepsilon^2-1\right)^2\left(k^2+\gamma^2\right)} - \frac{\varepsilon^2}{\left(\gamma^2\varepsilon^2-1\right)^2\left(\varepsilon^2k^2+1\right)} + \frac{\varepsilon^2}{\left(\gamma^2\varepsilon^2-1\right)\left(\varepsilon^2k^2+1\right)^2}, \tag{14}$$

and using more integral identities from the Appendix, we find

$$2A_\varepsilon(r) + B_\varepsilon(r) = \frac{1}{4\pi\left(\gamma^2\varepsilon^2-1\right)}\left[\frac{2\mathcal{K}_0(\gamma r) - 2\mathcal{K}_0(r/\varepsilon)}{\gamma^2\varepsilon^2-1} + \frac{r}{\varepsilon}\mathcal{K}_1(r/\varepsilon)\right], \tag{15a}$$

$$A_\varepsilon(r) + B_\varepsilon(r) = \frac{1}{2\pi r^2\gamma^2} - \frac{1}{4\pi\left(\gamma^2\varepsilon^2-1\right)}\left[\frac{2}{r\left(\gamma^2\varepsilon^2-1\right)}\left(\frac{\mathcal{K}_1(\gamma r)}{\gamma} - \varepsilon\mathcal{K}_1(r/\varepsilon)\right) + \mathcal{K}_2(r/\varepsilon)\right]. \tag{15b}$$

This is a system of simultaneous equations for $A_\varepsilon(r), B_\varepsilon(r)$, with solution

$$A_\varepsilon(r) = -\frac{1}{2\pi r^2\gamma^2} + \frac{1}{4\pi\left(\gamma^2\varepsilon^2-1\right)^2}\left[2\mathcal{K}_0(\gamma r) + \frac{2\mathcal{K}_1(\gamma r)}{\gamma r} + (\gamma^2\varepsilon^2-3)\mathcal{K}_0(r/\varepsilon) + \frac{2\gamma^2\varepsilon^4 + \left(\gamma^2r^2-4\right)\varepsilon^2 - r^2}{\varepsilon r}\mathcal{K}_1(r/\varepsilon)\right], \tag{16a}$$

$$B_\varepsilon(r)=\frac{1}{\pi r^2\gamma^2}-\frac{1}{4\pi\left(\gamma^2\varepsilon^2-1\right)^2}\left[2\left(\gamma^2\varepsilon^2-2\right)\mathcal{K}_0(r/\varepsilon)+\frac{4\gamma^2\varepsilon^4+\left(\gamma^2r^2-8\right)\varepsilon^2-r^2}{\varepsilon r}\mathcal{K}_1(r/\varepsilon)+2\mathcal{K}_2(\gamma r)\right], \tag{16b}$$

where we have used $\mathcal{K}_2(x)=\mathcal{K}_0(x)+2\mathcal{K}_1(x)/x$ [S12]. The singular limit $\varepsilon\gamma=1$ is regular: using Ref. [S12], we find

$$A_{\gamma^{-1}}(r)=-\frac{1}{2\pi r^2\gamma^2}+\frac{1}{16\pi}\left[\left(\gamma^2r^2+4\right)\mathcal{K}_0(\gamma r)+\left(3\gamma^2r^2+8\right)\frac{\mathcal{K}_1(\gamma r)}{\gamma r}\right], \tag{17a}$$

$$B_{\gamma^{-1}}(r)=\frac{1}{\pi r^2\gamma^2}-\frac{1}{16\pi}\left[\left(\gamma^2r^2+8\right)\mathcal{K}_0(\gamma r)+4\left(\gamma^2r^2+4\right)\frac{\mathcal{K}_1(\gamma r)}{\gamma r}\right]. \tag{17b}$$

This determines the Brinkmanlet (11) completely. The exponential decay of $\mathcal{K}_0,\mathcal{K}_1,\dots$ for $r\to\infty$ [S12] implies

$$\boldsymbol{u}\sim\frac{1}{2\pi r^2\gamma^2}\left(-\boldsymbol{f}+\frac{2(\boldsymbol{f}\cdot\boldsymbol{x})\boldsymbol{x}}{r^2}\right). \tag{18}$$

In particular, $\boldsymbol{u}\to\boldsymbol{0}$ as $r\to\infty$: There is no Stokes' paradox. Equation (18) also shows that, in the far field, $\boldsymbol{u}\parallel-\boldsymbol{f}$ in the direction $\boldsymbol{f}\cdot\boldsymbol{x}=0$. Clearly, $\boldsymbol{u}\parallel\boldsymbol{f}$ close to the origin (where the force is aplied the fluid), so this is a signature of lateral vortices in the flow field.

#### B. Solution of Eqs. (6): the regularised Brinkman sink

We solve Eqs. (6), again, in Fourier space. Taking their Fourier transform [S13], we find, for a radially symmetric blob function,

$$\boldsymbol{0}=-\mathrm{i}\boldsymbol{k}\tilde{p}-k^2\tilde{\boldsymbol{u}}-\gamma^2\tilde{\boldsymbol{u}},\qquad \mathrm{i}\boldsymbol{k}\cdot\tilde{\boldsymbol{u}}=-q\tilde{\phi}_\varepsilon(k). \tag{19}$$

Contracting the first equation with $\boldsymbol{k}$, using the second equation, and backsubstituting into the first now yields

$$\tilde{p}=-\left(1+\frac{\gamma^2}{k^2}\right)q\tilde{\phi}_\varepsilon(k)\quad\text{and}\quad\tilde{\boldsymbol{u}}=\frac{\mathrm{i}\boldsymbol{k}}{k^2}q\tilde{\phi}_\varepsilon(k). \tag{20}$$

By symmetry, the regularised Brinkman sink flow is

$$\boldsymbol{u}=C_\varepsilon(r)\boldsymbol{x}=\frac{1}{(2\pi)^2}\iint_{\mathbb{R}^2}\left(\frac{\mathrm{i}\boldsymbol{k}}{k^2}q\tilde{\phi}_\varepsilon(k)\right)\mathrm{e}^{\mathrm{i}\boldsymbol{k}\cdot\boldsymbol{x}}\,\mathrm{d}^2\boldsymbol{k}. \tag{21}$$

With Cartesian axes in which $\boldsymbol{x}=(r,0)$, $\boldsymbol{k}=k(\cos\theta,\sin\theta)$, the non-zero component of this relation yields, using the blob function in Eq. (8a),

$$C_\varepsilon(r)=\frac{1}{(2\pi)^2r}\int_0^\infty\int_0^{2\pi}\left(\mathrm{i}q\tilde{\phi}_\varepsilon(k)\cos\theta\right)\mathrm{e}^{\mathrm{i}kr\cos\theta}\,\mathrm{d}\theta\,\mathrm{d}k=\frac{-q}{2\pi r}\int_0^\infty\frac{\mathcal{J}_1(kr)}{\left(\varepsilon^2k^2+1\right)^2}\,\mathrm{d}k=\frac{-q}{2\pi r^2}\left[1-\frac{r^2}{2\varepsilon^2}\mathcal{K}_2(r/\varepsilon)\right], \tag{22}$$

where we have used integral identities from the Appendix. This determines the solution (21) of Eqs. (6) completely. We emphasise that the flow field is independent of the friction coefficient $\gamma$, though the pressure field depends on $\gamma$.

#### C. Fitting experimental flow fields

To fit the experimental flow field $\boldsymbol{U}(X,Y)$, where the $X,Y$ directions are left $\to$ right and vegetal $\to$ animal, respectively, we first determine the axis of mesendoderm migration $X=X_0$. We do so by minimising $\|\boldsymbol{U}(X,Y)-\boldsymbol{U}(2X_0-X,Y)\|$ with respect to $X_0$ using the MATLAB (The MathWorks, Inc.) function `fminsearch`, linearly interpolating the experimental flow fields onto a regular grid.

We then shift the coordinates so that $X_0=0$, and again interpolate $\boldsymbol{U}(X,Y)$ linearly on a regular grid $(X_i,Y_j)$ for $i,j=1,2,\dots,N=20$. We define $\boldsymbol{U_{i,j}}=\boldsymbol{U}(X_i,Y_j)$, and we decompose

$$\boldsymbol{U_{i,j}}=\overline{\boldsymbol{U}}_{\boldsymbol{i,j}}+\hat{\boldsymbol{U}}_{\boldsymbol{i,j}},\qquad\text{with }\overline{\boldsymbol{U}}_{\boldsymbol{i,j}}=\frac{\boldsymbol{U_{i,j}}+\boldsymbol{U_{N-i,j}}}{2}, \tag{23}$$

for $i,j=1,2,\dots,N$, so that $\overline{\boldsymbol{U}}_{\boldsymbol{i,j}}$ and $\hat{\boldsymbol{U}}_{\boldsymbol{i,j}}$ are, respectively, the left-right symmetric and asymmetric parts of the flow.

To fit the symmetric part of the experimental flow, we represent sinks and forces on and parallel to the axis of mesendoderm migration by a regularised Brinkman sink flow $\boldsymbol{u_0}$ of strength $q$ at $(0,y_0)$ and two regularised Brinkmanlet flows $\boldsymbol{u_1},\boldsymbol{u_2}$ resulting from forces $\boldsymbol{f_1}=(0,f_1)$, $\boldsymbol{f_2}=(0,f_2)$ at $(0,y_1)$, $(0,y_2)$, respectively. These regularised singularities result in a flow $\boldsymbol{u}=\boldsymbol{u_0}+\boldsymbol{u_1}+\boldsymbol{u_2}$, and we let $\boldsymbol{u_{i,j}}=\boldsymbol{u}(X_i,Y_j)$, for $i,j=1,2,\dots,N$. The fitting minimises the relative error

$$E=\sum_{i=1}^N\sum_{j=1}^N\left(\max\left\{\frac{\left\|\boldsymbol{u_{i,j}}-\overline{\boldsymbol{U}}_{\boldsymbol{i,j}}\right\|-\left\|\hat{\boldsymbol{U}}_{\boldsymbol{i,j}}\right\|}{\left\|\overline{\boldsymbol{U}}_{\boldsymbol{i,j}}\right\|},0\right\}\right)^2 \tag{24}$$

with respect to $y_0,y_1,y_2,q,f_1,f_2$, the friction parameter $\gamma$, and the regularisations $\varepsilon_0,\varepsilon_1=\varepsilon_2$ of the three singularities using `fminsearch` [S15]. Cutting off the relative error in this way ensures that differences between the fitted and observed flows below the asymmetry of the experimental data (as a proxy for experimental error) are not penalised. One could argue that minimising a relative rather than an absolute error puts too much weight on small flow velocities (the measurements of which may be unreliable), but we found that the fits minimising an absolute error tend to not represent topological features (e.g. vortices) of the experimental flows (not shown).

### III. EXTENSIONS OF THE MODEL

In this Section, we discuss extensions of the model to represent the YSL/EVL margin and the effects of compressibility.

#### A. Modelling the YSL/EVL margin

We represent the YSL/EVL margin as a boundary at $Y=Y_\mathrm{m}$, at which we apply the boundary condition $\boldsymbol{u}\cdot\hat{\boldsymbol{Y}}=0$, allowing flow parallel to, but not across the margin.

We add (regularised) image singularities to the flow: an image Brinkman sink flow $\boldsymbol{u}_0'$ of strength $q$ at $(0, 2Y_\mathrm{m} - y_0)$, and image Brinkmanlet flows $\boldsymbol{u}_1', \boldsymbol{u}_2'$ from forces $(0, -f_1), (0, -f_2)$ at the respective points $(0, 2Y_\mathrm{m} - y_1), (0, 2Y_\mathrm{m} - y_2)$. The resulting total flow $\boldsymbol{u} = \boldsymbol{u}_0 + \boldsymbol{u}_1 + \boldsymbol{u}_2 + \boldsymbol{u}_0' + \boldsymbol{u}_1' + \boldsymbol{u}_2'$ satisfies the boundary condition $\boldsymbol{u} \cdot \hat{\boldsymbol{Y}} = 0$ by symmetry. We emphasise that, given $Y_\mathrm{m}$, modelling the YSL/EVL margin does not add additional parameters to be fitted to the model.

#### B. Compressibility

A major simplifying assumption of our model is to assume the two-dimensional experimental flow to be incompressible: the cell sheet is essentially incompressible (since cells are mostly water), but in-plane compression of the cell sheet can be balanced by changes in its thickness, so incompressibility of the three-dimensional flow does not imply incompressibility of the two-dimensional flow. We do believe, however, that it is appropriate nevertheless to use an incompressible model since the compressive flows co-localise with the ingression of the cell sheet, so that the sink flow in the model can be interpreted as the sum of a contribution from compressibility of the flow, and the actual ingression flow.

Still, to discuss the effect of compressibility further, we derive here the expressions for (regularised) force singularities in a very compressible Brinkman flow. We start from the steady Cauchy equation without inertia [S16], with a friction force $-\gamma^2\boldsymbol{u}$ and a body force $\boldsymbol{F}(\boldsymbol{x})$,

$$\boldsymbol{0} = \boldsymbol{\nabla} \cdot \boldsymbol{\sigma} - \gamma^2 \boldsymbol{u} + \boldsymbol{F}(\boldsymbol{x}), \tag{25a}$$

where $\boldsymbol{\sigma}$ is the stress tensor. For a compressible fluid [S17],

$$\boldsymbol{\nabla} \cdot \boldsymbol{\sigma} = -\boldsymbol{\nabla} p + \nabla^2 \boldsymbol{u} + \alpha \boldsymbol{\nabla}(\boldsymbol{\nabla} \cdot \boldsymbol{u}), \tag{25b}$$

wherein $\alpha$ is the ratio of bulk to shear viscosity. The steady continuity equation with a sink $Q(\boldsymbol{x})$ is [S18]

$$\boldsymbol{\nabla} \cdot (\rho \boldsymbol{u}) = -Q(\boldsymbol{x}), \tag{25c}$$

in which $\rho(\boldsymbol{x})$ is the density of the fluid, which is coupled to the pressure via an equation of state $p = P(\rho)$ [S19]. Importantly, Eq. (25c) is nonlinear, so the solution of problem (25) cannot be expressed as a superposition of singularities.

Progress can however made in the particular case of a very compressible fluid with $p \approx 0$ [S19], in which case Eq. (25a) decouples from Eq. (25c) and becomes linear. The assumption $p \approx 0$ was previously made to describe the forces positioning the neural anlage in zebrafish [S20]. Here, we will thus seek (regularised) force singularities such that

$$\boldsymbol{0} = \nabla^2 \boldsymbol{u} + \alpha \boldsymbol{\nabla}(\boldsymbol{\nabla} \cdot \boldsymbol{u}) - \gamma^2 \boldsymbol{u} + \boldsymbol{f} \phi_\varepsilon(r). \tag{26}$$

We solve this equation, again, in Fourier space [S13]. We find, again assuming the blob function to be radially symmetric,

$$\boldsymbol{0} = -k^2 \tilde{\boldsymbol{u}} - \alpha \boldsymbol{k}(\boldsymbol{k} \cdot \tilde{\boldsymbol{u}}) - \gamma^2 \tilde{\boldsymbol{u}} + \boldsymbol{f} \tilde{\phi}_\varepsilon(k). \tag{27}$$

Contracting this equation with $\boldsymbol{k}$ yields

$$\boldsymbol{k} \cdot \tilde{\boldsymbol{u}} = \frac{\tilde{\phi}_\varepsilon(k)}{(1+\alpha)k^2 + \gamma^2} \boldsymbol{f} \cdot \boldsymbol{k}, \tag{28a}$$

whence

$$\tilde{\boldsymbol{u}} = \boldsymbol{f} \cdot \left[ \frac{\tilde{\phi}_\varepsilon(k)}{k^2 + \gamma^2} \left( \boldsymbol{I} - \frac{\alpha \boldsymbol{k}\boldsymbol{k}}{(1+\alpha)k^2 + \gamma^2} \right) \right]. \tag{28b}$$

The regularised compressible Brinkmanlet solution in two dimensions is therefore

$$\boldsymbol{u} = \boldsymbol{f} \cdot \left[ D_\varepsilon(r) \boldsymbol{I} + E_\varepsilon(r) \frac{\boldsymbol{x}\boldsymbol{x}}{r^2} \right] \equiv \boldsymbol{f} \cdot \boldsymbol{U}_\varepsilon(\boldsymbol{x}), \tag{29}$$

where the coefficients $D_\varepsilon(r), E_\varepsilon(r)$ are determined by the Fourier inversion integral

$$\boldsymbol{U}_\varepsilon(\boldsymbol{x}) = \frac{1}{(2\pi)^2} \iint_{\mathbb{R}^2} \frac{\tilde{\phi}_\varepsilon(k)}{k^2 + \gamma^2} \left( \boldsymbol{I} - \frac{\alpha \boldsymbol{k}\boldsymbol{k}}{(1+\alpha)k^2 + \gamma^2} \right) \mathrm{e}^{\mathrm{i}\boldsymbol{k}\cdot\boldsymbol{x}}\, \mathrm{d}^2\boldsymbol{k}. \tag{30}$$

Similarly to the derivation of Eqs. (13), it follows that

$$2D_\varepsilon(r) + E_\varepsilon(r) = \frac{1}{2\pi} \int_0^\infty \frac{k \tilde{\phi}_\varepsilon(k)}{k^2 + \gamma^2} \left[ 2 - \frac{\alpha k^2}{(1+\alpha)k^2 + \gamma^2} \right] \mathcal{J}_0(kr)\, \mathrm{d}k, \tag{31a}$$

$$D_\varepsilon(r) + E_\varepsilon(r) = \frac{1}{2\pi} \int_0^\infty \frac{k \tilde{\phi}_\varepsilon(k)}{k^2 + \gamma^2} \left\{ \left[ 1 - \frac{\alpha k^2}{(1+\alpha)k^2 + \gamma^2} \right] \mathcal{J}_0(kr) + \frac{\alpha k \mathcal{J}_1(kr)}{r\left[(1+\alpha)k^2 + \gamma^2\right]} \right\} \mathrm{d}k. \tag{31b}$$

For the blob function in Eq. (8a), using the partial fraction decompositions

$$\begin{aligned} \frac{\tilde{\phi}_\varepsilon(k)}{k^2 + \gamma^2} \left[ 2 - \frac{\alpha k^2}{(1+\alpha)k^2 + \gamma^2} \right] = {}& \frac{(1+\alpha)^2}{\left(\gamma^2\varepsilon^2 - 1 - \alpha\right)^2 \left[(1+\alpha)k^2 + \gamma^2\right]} + \frac{\varepsilon^2 \left(2\gamma^2\varepsilon^2 - 2 - \alpha\right)}{\left(\gamma^2\varepsilon^2 - 1\right)\left(\gamma^2\varepsilon^2 - 1 - \alpha\right)\left(\varepsilon^2 k^2 + 1\right)^2} \\ & + \frac{1}{\left(\gamma^2\varepsilon^2 - 1\right)^2 \left(k^2 + \gamma^2\right)} - \frac{\varepsilon^2 \left[(1+\alpha)(2+\alpha) - 4(1+\alpha)\gamma^2\varepsilon^2 + (\alpha+2)\gamma^4\varepsilon^4\right]}{\left(\gamma^2\varepsilon^2 - 1\right)^2 \left(\gamma^2\varepsilon^2 - 1 - \alpha\right)^2 \left(\varepsilon^2 k^2 + 1\right)}, \end{aligned} \tag{32a}$$

$$\begin{aligned} \frac{\tilde{\phi}_\varepsilon(k)}{k^2 + \gamma^2} \left[ 1 - \frac{\alpha k^2}{(1+\alpha)k^2 + \gamma^2} \right] = {}& \frac{(1+\alpha)^2}{\left(\gamma^2\varepsilon^2 - 1 - \alpha\right)^2 \left[(1+\alpha)k^2 + \gamma^2\right]} - \frac{(1+\alpha)\varepsilon^2}{\left(\gamma^2\varepsilon^2 - 1 - \alpha\right)^2 \left(\varepsilon^2 k^2 + 1\right)} \\ & + \frac{\varepsilon^2}{\left(\gamma^2\varepsilon^2 - 1 - \alpha\right)\left(\varepsilon^2 k^2 + 1\right)^2}, \end{aligned} \tag{32b}$$

$$\frac{\alpha k^2\tilde{\phi}_\varepsilon(k)}{(k^2+\gamma^2)\left[(1+\alpha)k^2+\gamma^2\right]} = \frac{1}{\left(\gamma^2\varepsilon^2-1\right)^2(k^2+\gamma^2)} - \frac{(1+\alpha)^2}{\left(\gamma^2\varepsilon^2-1-\alpha\right)^2\left[(1+\alpha)k^2+\gamma^2\right]} - \frac{\alpha\varepsilon^2}{\left(\gamma^2\varepsilon^2-1\right)\left(\gamma^2\varepsilon^2-1-\alpha\right)\left(\varepsilon^2k^2+1\right)^2} + \frac{\alpha\varepsilon^2\left(\gamma^4\varepsilon^4-1-\alpha\right)}{\left(\gamma^2\varepsilon^2-1\right)^2\left(\gamma^2\varepsilon^2-1-\alpha\right)^2\left(\varepsilon^2k^2+1\right)}, \tag{32c}$$

and integrals from the Appendix, we obtain

$$2D_\varepsilon(r)+E_\varepsilon(r) = \frac{\mathcal{K}_0(\gamma r)}{2\pi\left(\gamma^2\varepsilon^2-1\right)^2} + \frac{(1+\alpha)\mathcal{K}_0\left(\frac{\gamma r}{\sqrt{1+\alpha}}\right)}{2\pi\left(\gamma^2\varepsilon^2-1-\alpha\right)^2} - \frac{\left[(1+\alpha)(2+\alpha)-4(1+\alpha)\gamma^2\varepsilon^2+(\alpha+2)\gamma^4\varepsilon^4\right]\mathcal{K}_0(r/\varepsilon)}{2\pi\left(\gamma^2\varepsilon^2-1\right)^2\left(\gamma^2\varepsilon^2-1-\alpha\right)^2} + \frac{\left(2\gamma^2\varepsilon^2-2-\alpha\right)r\mathcal{K}_1(r/\varepsilon)}{4\pi\varepsilon\left(\gamma^2\varepsilon^2-1\right)\left(\gamma^2\varepsilon^2-1-\alpha\right)}, \tag{33a}$$

$$D_\varepsilon(r)+E_\varepsilon(r) = \frac{(1+\alpha)\mathcal{K}_0\left(\frac{\gamma r}{\sqrt{1+\alpha}}\right)}{2\pi\left(\gamma^2\varepsilon^2-1-\alpha\right)^2} - \frac{\left[(\alpha+2)\gamma^2\varepsilon^2+(\alpha+1)(\alpha-2)\right]\mathcal{K}_0(r/\varepsilon)}{2\pi\left(\gamma^2\varepsilon^2-1\right)\left(\gamma^2\varepsilon^2-1-\alpha\right)^2} - \frac{\mathcal{K}_1(\gamma r)}{2\pi\gamma r\left(\gamma^2\varepsilon^2-1\right)^2} + \frac{(1+\alpha)^{3/2}\mathcal{K}_1\left(\frac{\gamma r}{\sqrt{1+\alpha}}\right)}{2\pi\gamma r\left(\gamma^2\varepsilon^2-1-\alpha\right)^2} + \left\{\frac{\alpha\varepsilon\left[2(1+\alpha)-(2+\alpha)\gamma^2\varepsilon^2\right]}{r\left(\gamma^2\varepsilon^2-1-\alpha\right)\left(\gamma^2\varepsilon^2-1\right)^2} + \frac{r}{2\varepsilon}\right\}\frac{\mathcal{K}_1(r/\varepsilon)}{2\pi\left(\gamma^2\varepsilon^2-1-\alpha\right)}, \tag{33b}$$

where, again, we have used $\mathcal{K}_2(x) = \mathcal{K}_0(x) + 2\mathcal{K}_1(x)/x$ [S12]. Thence

$$D_\varepsilon(r) = \frac{\mathcal{K}_0(\gamma r)}{2\pi\left(\gamma^2\varepsilon^2-1\right)^2} - \frac{\left[(2+\alpha)\gamma^2\varepsilon^2-2-3\alpha\right]\mathcal{K}_0(r/\varepsilon)}{4\pi\left(\gamma^2\varepsilon^2-1-\alpha\right)\left(\gamma^2\varepsilon^2-1\right)^2} + \frac{\mathcal{K}_1(\gamma r)}{2\pi\gamma r\left(\gamma^2\varepsilon^2-1\right)^2} - \frac{(1+\alpha)^{3/2}\mathcal{K}_1\left(\frac{\gamma r}{\sqrt{1+\alpha}}\right)}{2\pi\gamma r\left(\gamma^2\varepsilon^2-1-\alpha\right)^2} + \left\{\frac{\alpha\varepsilon\left[(2+\alpha)\gamma^2\varepsilon^2-2(1+\alpha)\right]}{r\left(\gamma^2\varepsilon^2-1-\alpha\right)^2\left(\gamma^2\varepsilon^2-1\right)} + \frac{r}{2\varepsilon}\right\}\frac{\mathcal{K}_1(r/\varepsilon)}{2\pi\left(\gamma^2\varepsilon^2-1\right)}, \tag{34a}$$

$$E_\varepsilon(r) = \frac{\alpha\left[2(1+\alpha)-(2+\alpha)\gamma^2\varepsilon^2\right]\mathcal{K}_0(r/\varepsilon)}{2\pi\left(\gamma^2\varepsilon^2-1-\alpha\right)^2\left(\gamma^2\varepsilon^2-1\right)^2} + \left\{\frac{\varepsilon\left[4(1+\alpha)-2(2+\alpha)\gamma^2\varepsilon^2\right]}{r\left(\gamma^2\varepsilon^2-1-\alpha\right)\left(\gamma^2\varepsilon^2-1\right)} + \frac{r}{2\varepsilon}\right\}\frac{\alpha\mathcal{K}_1(r/\varepsilon)}{2\pi\left(\gamma^2\varepsilon^2-1-\alpha\right)\left(\gamma^2\varepsilon^2-1\right)} - \frac{\mathcal{K}_2(\gamma r)}{2\pi\left(\gamma^2\varepsilon^2-1\right)^2} + \frac{(1+\alpha)\mathcal{K}_2\left(\frac{\gamma r}{\sqrt{1+\alpha}}\right)}{2\pi\left(\gamma^2\varepsilon^2-1-\alpha\right)^2}. \tag{34b}$$

The singular limits $\varepsilon\gamma = 1$ and $\varepsilon\gamma = \sqrt{1+\alpha}$ are, again, regular, and we find, using Ref. [S12],

$$D_{\gamma^{-1}}(r) = \frac{1}{16\pi\alpha^2}\left\{\left[4\alpha(1+\alpha)+\alpha^2r^2\gamma^2\right]\mathcal{K}_0(\gamma r) + \left[\frac{8(1+\alpha)^2}{\gamma r}+3\alpha^2\gamma r\right]\mathcal{K}_1(\gamma r) - \frac{8(1+\alpha)^{3/2}}{\gamma r}\mathcal{K}_1\left(\frac{\gamma r}{\sqrt{1+\alpha}}\right)\right\}, \tag{35a}$$

$$E_{\gamma^{-1}}(r) = \frac{1}{16\pi\alpha^2}\left\{-\left[8(1+\alpha)^2+\alpha^2r^2\gamma^2\right]\mathcal{K}_0(\gamma r) - 4(1+\alpha)\left[4(1+\alpha)+\alpha\gamma^2r^2\right]\frac{\mathcal{K}_1(\gamma r)}{\gamma r} + 8(1+\alpha)\mathcal{K}_2\left(\frac{\gamma r}{\sqrt{1+\alpha}}\right)\right\}, \tag{35b}$$

$$D_{\frac{\sqrt{1+\alpha}}{\gamma}}(r) = \frac{1}{16\pi\alpha^2}\left[8\mathcal{K}_0(\gamma r) + \frac{8\mathcal{K}_1(\gamma r)}{\gamma r} - \frac{4(2+\alpha)}{1+\alpha}\mathcal{K}_0\left(\frac{\gamma r}{\sqrt{1+\alpha}}\right) + \frac{\alpha(4+3\alpha)\gamma^2r^2-8(1+\alpha)}{(1+\alpha)^{3/2}\gamma r}\mathcal{K}_1\left(\frac{\gamma r}{\sqrt{1+\alpha}}\right)\right], \tag{35c}$$

$$E_{\frac{\sqrt{1+\alpha}}{\gamma}}(r) = \frac{1}{16\pi\alpha^2}\left\{-8\mathcal{K}_2(\gamma r) + \frac{8(1+\alpha)+\alpha^2\gamma^2r^2}{(1+\alpha)^2}\mathcal{K}_0\left(\frac{\gamma r}{\sqrt{1+\alpha}}\right) + \frac{4\left[4(1+\alpha)-\alpha\gamma^2r^2\right]}{(1+\alpha)^{3/2}\gamma r}\mathcal{K}_1\left(\frac{\gamma r}{\sqrt{1+\alpha}}\right)\right\}. \tag{35d}$$

This determines the compressible Brinkmanlet (29) completely. We next consider the behaviour as $r \longrightarrow \infty$ in the direction $\boldsymbol{f}\cdot\boldsymbol{x} = 0$ perpendicular to the applied force. From the asymptotic behaviour as $r \longrightarrow \infty$ of $\mathcal{K}_0, \mathcal{K}_1, \mathcal{K}_2$ [S12], we find

$$\boldsymbol{u} \overset{\boldsymbol{f}\cdot\boldsymbol{x}=0}{\sim} \frac{\boldsymbol{f}}{\sqrt{8\pi}}\left\{e^{-\frac{\gamma r}{\sqrt{1+\alpha}}}\left[-\frac{(1+\alpha)^{7/4}r^{-3/2}}{\gamma^{3/2}\left(\gamma^2\varepsilon^2-1-\alpha\right)^2}+\cdots\right] + e^{-r/\varepsilon}\left[\frac{r^{1/2}}{2\varepsilon^{1/2}\left(\gamma^2\varepsilon^2-1\right)}+\cdots\right]\right\}, \tag{36}$$

where we have used the fact that $e^{-\gamma r} \ll e^{-\frac{\gamma r}{\sqrt{1+\alpha}}}$ for $\alpha > 0$. Hence there are two cases: if $\gamma\varepsilon < \sqrt{1+\alpha}$, then the first term dominates in the far-field and $\boldsymbol{u}\cdot\boldsymbol{f} < 0$. This is again the far-field signature of lateral vortices in the flow field. If however $\gamma\varepsilon > \sqrt{1+\alpha} > 1$, then the second term dominates, and $\boldsymbol{u}\cdot\boldsymbol{f} > 0$, which suggests that there are no lateral vortices

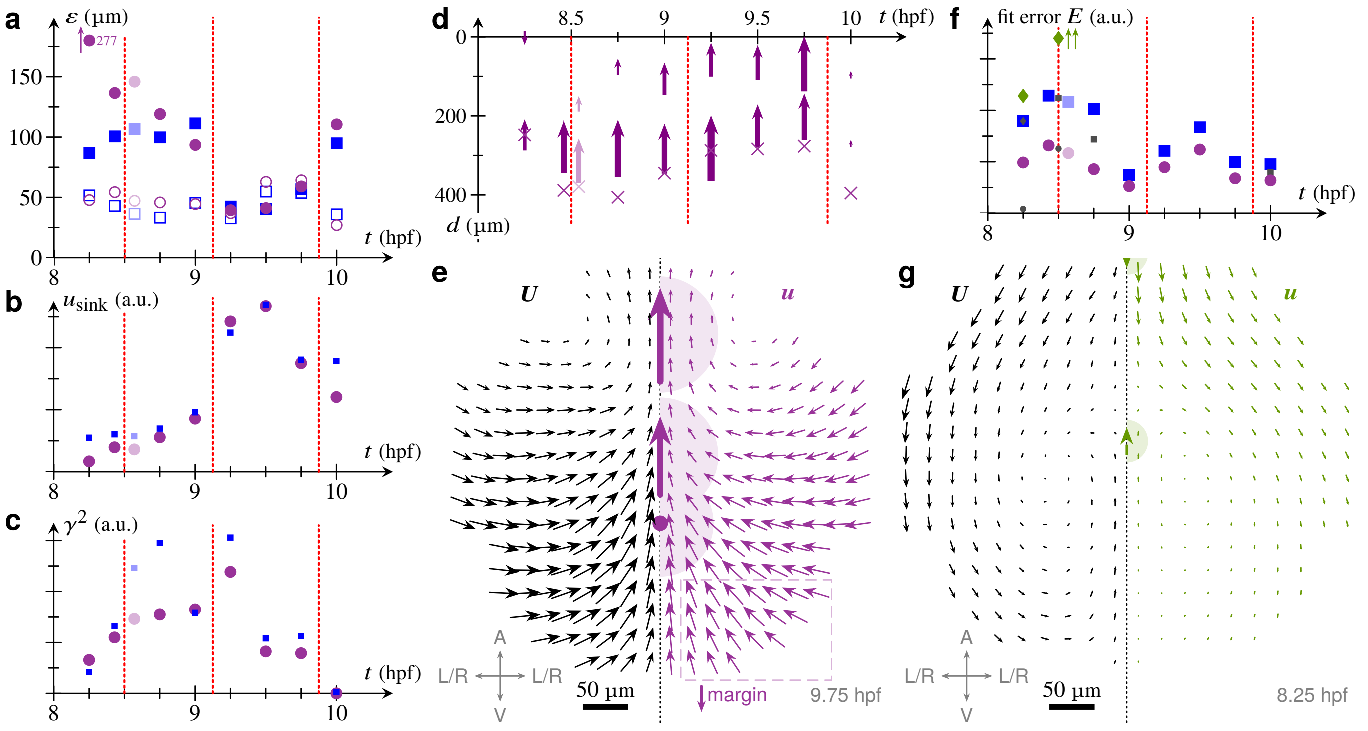


FIG. S1. Extensions of the model. (a) Fitted values of the sink regularisation radii $\varepsilon_0$ (filled markers) and force regularisation radii $\varepsilon_1 = \varepsilon_2$ (open markers), as a function of time $t$, expressed in hours-post-fertilisation (hpf). Results are shown for the model without margin (square markers) and the model with margin (round markers). Dashed vertical lines represent the three transitions discussed in the main text. Where two qualitatively different physical fits were found for one timepoint (close to one of these transitions), values for both fits are shown in a staggered fashion, with the temporal order decided by the other fits and the mechanical and biological interpretation of the transition. An offscale value is indicated by a vertical arrow. (b) Fitted fluid velocity $u_{\text{sink}}$, in arbitrary units (a.u.), induced by the sink at the position of the sink as a function of time $t$. Round markers show results for the model with margin; small square markers repeat the results for the model without margin from Fig. 3n of the main text for comparison. (c) Analogous plot of the fitted friction coefficient $\gamma^2$ against time $t$. Results for the model without margin are repeated from Fig. 3o of the main text. (d) Plot of the positions of the fitted regularised sinks (crosses) and of the positions, orientations, and magnitudes of the fitted regularised Brinkmanlets (arrows) as a function of time $t$, for the model with margin, analogous to Fig. 3m of the main text. The positions are expressed in terms of the distance $d$ from the anterior end of the ANP. The image singuarities imposing the margin boundary condition are not shown. (e) Left-right symmetrised experimental flow field $\boldsymbol{U}$ (left), reproduced by the fitted flow field $\boldsymbol{u}$ (right) for the model with margin at $t = 9.75$ hpf, analogous to Fig. 3l of the main text. Arrows on the axis represent the positions, directions, and magnitudes of the fitted regularised Brinkmanlets; the dot represents the position of the fitted regularised Brinkman sinks. Shaded areas show the corresponding regularisation radii. The image singularities are, again, not shown. The dashed box highlights the underestimate of the left-right flows by the fit. The arrow and "margin" label emphasise the vegetal or posterior direction towards the boundary at which the margin boundary condition is applied. Inset: embryo axes (A: animal, V: vegetal; L/R: left-right symmetrised). (f) Fit error $E$, defined by Eq. (24), as a funtion of time $t$ for the model without margin (square markers), the model with margin (round markers), and the infinitely compressible model (lozenge markers). Small markers show lower fit errors for discarded, unphysical fits for some timepoints. Double vertical arrows indicate an offscale value. (g) Experimental and fitted flow fields for the infinitely compressible model at $t = 8.25$ hpf, analogous to Fig. 3k of the main text.

in this case. This suggestion is confirmed by numerical calculations (not shown). This shows that, for the compressible Brinkmanlet, changes in $\varepsilon$ can change the topology of the flow field; this is not the case for the incompressible Brinkmanlet discussed earlier, for which the far-field behaviour, expressed by Eq. (18), is independent of $\varepsilon$.

There is no analogous problem for a regularised sink flow; indeed, with $p = 0$ and $\boldsymbol{F} = \boldsymbol{0}$, the solution of Eq. (25a) decaying in the far-field is simply $\boldsymbol{u} = \boldsymbol{0}$. The mass conservation equation (25c) then constitutes a contradiction. This is the mathematical reason why we use a model of an incompressible fluid to represent the experimental flows and in particular the sink resulting from ingression.

#### C. Discussion

Results for these extensions of the model, as well as for the fitted regularisation radii not discussed in the main text, are shown in Fig. S1. We discuss these results and their physical and biological implications below.

##### 1. *Fitted regularisation radii*

The fitted values of the regularisation radii allow for a consistency check of the fit: The force regularisation radii $\varepsilon_1 = \varepsilon_2$ represent the extent over which the mesendoderm exerts friction on the neurectoderm, so are expected not to change over time. Consistently with this expectation, the fitted values of $\varepsilon_1 = \varepsilon_2$ are roughly constant (Fig. S1a). By constrast, the fitted sink radius $\varepsilon_0$ decreases, as expected, as ingression movements become stronger (Fig. S1a): at this time, the sink represents localised ingression (small $\varepsilon_0$), but it represents the smeared-out, effective compressibility of the tissue (larger $\varepsilon_0$) at earlier times before ingression, due the assumption of incompressibility of the fluid. This interpretation is also consistent with the non-zero sink velocities fitted at the earliest timepoints (Fig. S1b), before the start of internalisation.

#### 2. *The EVL/YSL margin*

By representing the tissue as an infinite two-dimensional fluid, the model neglects the boundary conditions at the EVL/YSL margin. Because the margin is closest to the ANP in the vegetal direction, we represent the EVL/YSL margin as a line boundary at $Y = Y_\mathrm{m} < 0$, at which we impose a boundary condition of no flow across the boundary, as described above. These boundary position and condition are, of course, still a simplification that neglects the flows arising from continued epiboly movements. In particular, this new boundary condition might be expected to strengthen left-right flows at the vegetal end of the ANP, which are underestimated by the model without this boundary condition (Fig. 3l in the main text). [This underestimate might also result, however and *a priori*, from the fact that the uncertainty in the experimental flow field (i.e. its left-right asymmetry) is larger there (not shown).] For fitting the model with this new boundary condition, we fix $Y_\mathrm{m}$ so that the boundary is approximately 800 µm behind the front of the ANP. This value corresponds to the margin position at 7.5 hpf. We make this choice to maximise the effect of the boundary condition even though, because of continued epiboly movements, the margin will be farther from the ANP at later stages.

The resulting fit (Fig. S1a–e) agrees with the fit of the model without the margin. In particular, it recovers the three mechanical regimes discussed in the main text (Fig. S1b–d) and the fitted regularisation radii (Fig. S1a) continue to be consistent with expectations. Including the margin does not improve the fit of the vegetal left-right flows at late stages (Fig. S1e), though, suggesting that these discrepancies are due to the larger uncertainty of the experimental flows there. Finally, the fit error (Fig. S1f) is decreased compared to the case without the margin (without additional fitting parameters, since $Y_\mathrm{m}$ is not fitted for). This suggests that the simplified margin boundary condition correctly captures additional physics of the system.

#### 3. *Incompressibility*

As noted above, the assumption of an incompressible fluid implies that the compressibility of the tissue must be absorbed, at least for early stages, into an effective sink. An alternative description of the early stages assumes, as described earlier, an infinitely compressible fluid. Fitting only two regularised compressible Stokeslets representing the opposing forces from the mesendoderm, this model can represent the experimental flows at early stages quantitatively (Fig. S1g), but even at 8.25 hpf, the fit error is higher than that of the incompressible models with or without margin (Fig. S1f), and is much higher than that of the incompressible models from 8.5 hpf (Fig. S1f). (Note however that the compressible model has fewer fitting parameters.) This hints, physically, that a regularised sink in an incompressible fluid is a better mechanical representation of the compressibility of the system than an infinitely compressible fluid. It also hints, more biologically, that ingression movements (corresponding to a true sink, rather than an effective one) start to contribute to the experimental flows from 8.5 hpf, i.e. from the time of the first mechanical transition, when mesendoderm migration ceases to dominate the mechanics.

#### 4. *Ingression as a sink*

We conclude by briefly discussing our modelling of ingression as a sink. We note that we rejected some fits as being unphysical, although such fits can be found at lower fit scores than the fits that we have retained (Fig. S1g). We declared some such fits to be unphysical because, for example, the fitted values of the regularisation radii were much larger than the physical extent of the neurectoderm. The existence of such spurious fits is not surprising given the considerable nonlinearity of the fitting problem.

More interestingly, especially at late stages (> 9 hpf), we found fits in which (almost) equal and opposite forces are nearly colocalised with the sink (not shown). While colocalisation of the two fitted forces contradicts our physical interpretation of these forces (as forces exerted on the neurectoderm by the underlying mesendoderm), this may signify that the ingression flows are better represented by the superposition of a sink and a Brinkmanlet dipole. Fully understanding the physics of the ingression flows remains a question for future work. Meanwhile, extending our fitting procedure to fit two forces and such a superposition of a sink and a Brinkmanlet dipole is beyond the scope of our analysis, the more so as we cannot justify fitting four singularities in the same way as we justified the fitting of three singularities based on our argument showing that they are the minimal ingredients that can reproduce the flow fields even qualitatively (Fig. 3a–j of the main text, and discussion thereof).

### APPENDIX: INTEGRALS

In this Appendix, we give several integral identities required for the derivation of regularised singularities in this Supplemental Note. These identities can be obtained using Mathematica (Wolfram, Inc.).

Calculation of polar integrals in two-dimensional Fourier inversions requires the identities

$$\int_0^{2\pi} \mathrm{e}^{\mathrm{i}a\cos\theta}\,\mathrm{d}\theta = 2\pi\mathcal{J}_0(a), \tag{A1a}$$

$$\int_0^{2\pi} \mathrm{e}^{\mathrm{i}a\cos\theta}\cos\theta\,\mathrm{d}\theta = 2\mathrm{i}\pi\mathcal{J}_1(a), \tag{A1b}$$

$$\int_0^{2\pi} \mathrm{e}^{\mathrm{i}a\cos\theta}\sin^2\theta\,\mathrm{d}\theta = \frac{2\pi\mathcal{J}_1(a)}{a}, \tag{A1c}$$

valid for $a > 0$ and wherein $\mathcal{J}_0, \mathcal{J}_1, \dots$ again denote the Bessel functions of the first kind.

The calculations in this Supplemental Note further rely on the identities

$$\int_0^\infty \frac{\ell \mathcal{J}_0(a\ell)}{1+\ell^2}\,\mathrm{d}\ell = \mathcal{K}_0(a\ell), \tag{A2a}$$

$$\int_0^\infty \frac{\mathcal{J}_1(a\ell)}{1+\ell^2}\,\mathrm{d}\ell = \frac{1}{a} - \mathcal{K}_1(a\ell), \tag{A2b}$$

$$\int_0^\infty \frac{\ell \mathcal{J}_0(a\ell)}{\left(1+\ell^2\right)^2}\,\mathrm{d}\ell = \frac{a\mathcal{K}_1(a\ell)}{2}, \tag{A2c}$$

$$\int_0^\infty \frac{\mathcal{J}_1(a\ell)}{\left(1+\ell^2\right)^2}\,\mathrm{d}\ell = \frac{1}{a} - \frac{a\mathcal{K}_2(a\ell)}{2}, \tag{A2d}$$

valid again for $a > 0$, and in which $\mathcal{K}_0, \mathcal{K}_1, \mathcal{K}_2, \dots$ denote again the modified Bessel functions of the second kind.


[S1] Cæsar, *Commentarii de bello Gallico* (Ancient Rome); for a modern translation, see, e.g., *The Gallic War* (Penguin Classics, London, UK, 1982).

[S2] C. Pozrikidis, *Introduction to Theoretical and Computational Fluid Dynamics*, 2nd ed. (Oxford University Press, New York, NY, 2011) Chaps. 2.1, 6.1, 6.5, 6.6, pp. 112–123, 370–381, 410–431.

[S3] M. A. Goldshtik, Viscous-flow paradoxes, Ann. Rev. Fluid Mech. **22**, 441 (1990).

[S4] More mathematically, these anterior or posterior stagnation points disappear if Stokes' paradox is solved by the addition of a friction force $-\gamma^2\boldsymbol{u}$ to the flow as in Sec. II. Indeed, taking the limit $\varepsilon \longrightarrow 0$ in Eqs. (16) using Mathematica, the flow along the axis defined by $\boldsymbol{f}$ is, from Eq. (11), $\boldsymbol{u} = [A_0(r) + B_0(r)]\boldsymbol{f}$, with $A_0(r) + B_0(r) = [1 - \gamma r\mathcal{K}_1(\gamma r)]/(2\pi r^2\gamma^2) > 0$, since $x\mathcal{K}_1(x) < 1$ for $x > 0$. This shows, mathematically, that there are no stagnation points on the flow axis.

[S5] H. C. Brinkman, A calculation of the viscous force exerted by a flowing fluid on a dense swarm of particles, Flow Turb. Combust. **1**, 27 (1949).

[S6] R. Cortez, The method of regularized Stokeslets, SIAM J. Sci. Comput. **23**, 1204 (2001).

[S7] R. Cortez, L. Fauci, and A. Medovikov, The method of regularized Stokeslets in three dimensions: Analysis, validation, and application to helical swimming, Phys. Fluids **17**, 031504 (2005).

[S8] D. J. Smith, A nearest-neighbour discretisation of the regularized stokeslet boundary integral equation, J. Comp. Phys. **358**, 88 (2018).

[S9] C. Pozrikidis, *Boundary Integral and Singularity Methods for Linearized Viscous Flow*, Cambridge Texts in Applied Mathematics (Cambridge University Press, Cambridge, UK, 1992).

[S10] B. Zhao, E. Lauga, and L. Koens, Method of regularized stokeslets: Flow analysis and improvement of convergence, Phys. Rev. Fluids **4**, 084104 (2019).

[S11] K. Leiderman and S. D. Olson, Swimming in a two-dimensional Brinkman fluid: Computational modeling and regularized solutions, Phys. Fluids **28**, 021902 (2016).

[S12] M. Abramowitz and I. A. Stegun, *Handbook of Mathematical Functions*, 10th ed., Applied Mathematics Series, Vol. 55 (National Bureau of Standards, 1972) Chap. 9, pp. 355–389.

[S13] Throughout this Supplemental Note, we adopt the convention that the Fourier transform $\tilde{f}$ of a suitable function $f$ on $\mathbb{R}^2$ is

$$\tilde{f}(\boldsymbol{k}) = \iint_{\mathbb{R}^2} f(\boldsymbol{x})\mathrm{e}^{-\mathrm{i}\boldsymbol{k}\cdot\boldsymbol{x}}\,\mathrm{d}^2\boldsymbol{x}.$$

With this convention,

$$f(\boldsymbol{x}) = \frac{1}{(2\pi)^2}\iint_{\mathbb{R}^2} \tilde{f}(\boldsymbol{k})\mathrm{e}^{\mathrm{i}\boldsymbol{k}\cdot\boldsymbol{x}}\,\mathrm{d}^2\boldsymbol{k}$$

is the form taken by the Fourier inversion theorem in two dimensions.

[S14] Mathematica (Wolfram, Inc.) was used for the calculations in the Supplemental Note to assist with manipulating complicated algebraic expressions.

[S15] In more detail, we fit the experimental data in two steps: first, for each stage, we fit $y_0, y_1, y_2, q, f_1, f_2$ at fixed values of $\gamma, \varepsilon_0, \varepsilon_1 = \varepsilon_2$. At this stage, we also fit separately for different signs of $f_1, f_2$, and different relative positions of the singularities. We then seek local minima of $E$ on the grid of $\gamma, \varepsilon_0, \varepsilon_1 = \varepsilon_2$, and use these as initial guesses for a second fitting round with respect to all parameters. This enables us to find local minima in the fitting landscape, and hence discard unphysical fits resulting from unphysical global minima for some timepoints (as discussed further in Sec. III C of this Supplemental Note).

[S16] D. J. Acheson, *Elementary Fluid Dynamics*, Oxford Applied Mathematics and Computer Sciences Series (Clarendon Press, Oxford, UK, 2009) Chap. 6.3, pp. 205–207.

[S17] I. G. Currie, *Fundamental Mechanics of Fluids*, 2nd ed. (McGraw-Hill, New York, NY, 1993) Chaps. 1.11, 1.12, 1.13, pp. 23–29.

[S18] G. K. Batchelor, *An Introduction to Fluid Dynamics*, Cambridge Mathematical Library (Cambridge University Press, Cambridge, UK, 2000) Chap. 3.1, pp. 131–137.

[S19] For a one-dimensional model of a biological tissue, P. Recho, J. Ranft, and P. Marcq, One-dimensional collective migration of a proliferating cell monolayer, Soft Matter **12**, 2381 (2016) obtain $P(\rho) \sim -P_0 \log(\rho/\rho_0)$, where $P_0 = \ell/\tau$, with $\ell$ an Onsager coefficient and $\tau$ the characteristic time scale of cell division. In particular, $p \approx 0$ holds for very slow cell divisions.

[S20] M. Smutny, Z. Ákos, S. Grigolon, S. Shamipour, V. Ruprecht, D. Čapek, M. Behrndt, E. Papusheva, M. Tada, B. Hof, T. Vicsek, G. Salbreux, and C.-P. Heisenberg, Friction forces position the neural anlage, Nat. Cell Biol. **19**, 306 (2017).
